# In vivo reprogramming and epigenetic rejuvenation of adult cardiomyocytes ameliorate heart failure in mice

**DOI:** 10.1101/2021.12.22.473302

**Authors:** Irene de Lázaro, Bohan Zhang, Nadezhda E Makarova, Marco Mariotti, Tiara L Orejón-Sánchez, Christina M Tringides, Vadim N Gladyshev, David J Mooney

## Abstract

Partial cell reprogramming has been demonstrated in certain mouse tissues by in situ overexpression of *Oct3/4*, *Klf4*, *Sox2* and *cMyc* (OKSM) transcription factors, and can induce rejuvenation and/or augment regeneration. Reprogramming of adult cardiomyocytes has been elusive until recently, but its success could help overcome the lack of endogenous regenerative capacity of the mammalian myocardium. Here, we generated cardiomyocyte-specific, doxycycline-inducible, reprogrammable mice and demonstrated that sustained OKSM induction reprograms cardiomyocytes fully into teratoma-forming pluripotent cells. However, we also showed that cyclic OKSM upregulation induces significant decrease of epigenetic age in the cardiomyocytes without de-differentiation or reacquisition of pluripotency. In mice with progressive heart failure, cardiomyocyte epigenetic rejuvenation correlated with stabilization of systolic heart function. These findings confirm that OKSM can reprogram adult mouse cardiomyocytes to different states depending on the duration of their expression, and provide further evidence that partially reprogrammed cardiomyocytes may contribute to ameliorate cardiac disease.

## INTRODUCTION

Expression of *Oct3/4*, *Klf4*, *Sox2* and *cMyc* (OKSM) transcription factors in adult mouse tissues generates reprogrammed cells in situ^1–3^. These commit to the pluripotent state and can form undifferentiated tumors, or teratomas, when OKSM expression is sustained for extended periods^2^. However, reprogramming is a stepwise process^4^ and, with shorter duration of OKSM expression, cells undergo *partial* reprogramming. This is a so far broadly-defined term that can range from epigenetic rejuvenation – the erasure of epigenetic marks associated with cellular ageing^5, 6^ – to transient and reversible loss of cell identity, accompanied by time-limited proliferation and reactivation of pluripotency-related genes^1, 3^. Partial *in vivo* cell reprogramming has been proposed as an approach to enhance the regenerative capabilities of adult tissues that could benefit from such features as the lack of *ex vivo* manipulations, the versatility of OKSM factors to enable reprogramming of a wide variety of cell types, and the typically enhanced cell maturation that is achieved *in vivo* versus *in vitro*^7, 8^. To date, preclinical studies in mice have demonstrated that *in vivo* partial reprogramming through OKSM overexpression enhances regeneration of skeletal muscle^3, 9^, retina^6^ and liver^10^, improves wound healing^11^ and rejuvenates several tissues in progeroid and naturally aged mice^5, 6, 12, 13^. A recent study has also suggested that reprogramming early in life could have long-lasting effects, significantly increasing life-span and overall fitness at old age^14^. Confirmation of the regenerative and rejuvenating effects of *in vivo* partial reprogramming in multiple independent laboratories, in different tissues and disease models, supports the promise of this strategy.

However, most studies have focused on demonstrating improvements linked to cell reprogramming in tissues and organs without identifying the specific cell type (or types) that undergo this process^3, 5, 11, 14^. Many utilize transgenic *reprogrammable* mice that exploit tetracycline-inducible systems to control OKSM expression^2, 5, 11, 14^. While this strategy allows high levels of OKSM expression that ensure efficient reprogramming and tight temporal control over reprogramming, required to avoid tumorigenesis, OKSM expression is typically ubiquitous and thus does not enable cell-type specificity of reprogramming. Delivery of exogenous OKSM factors, with or without viral vectors, can offer tissue targeting capabilities but typically not with cell type resolution and is often limited by poor temporal control of OKSM expression. For example, integrating retroviruses and adeno-associated viruses (AAV) trigger tumorigenesis due to extended OKSM expression^15, 16^ while plasmid DNA and non-integrating adenoviruses have shown limited efficiency of reprogramming^3, 17^.

Given the negligible turnover of adult mammalian cardiomyocytes^18^, which severely limits the regenerative capacity of the heart, testing the possibility of *in vivo* partial cardiomyocyte reprogramming has been of particular interest. However, the ability of terminally differentiated, adult cardiomyocytes to undergo OKSM-induced reprogramming has been unclear. A negative correlation between a cell’s differentiation state and its capability to undergo OKSM-triggered reprogramming has been consistently described for various cell lineages^19–21^, which may explain why reprogramming of cardiac tissues has not been reported in reprogrammable mice with ubiquitous OKSM expression even after they developed lethal teratomas in other organs (mainly the pancreas and gut)^2^. Adenoviral vectors have been shown to induce transient reprogramming in neonatal rat and mouse cardiomyocytes *in vitro*^22^, but the same effect has not been confirmed in adult counterparts *in vivo*^17^. Only recently, Chen et al have demonstrated reprogramming-induced cell cycle re-entry of adult mouse cardiomyocytes, which improved post-ischemia myocardial regeneration and was achieved by relying on the inducible expression of OKSM under the control of a cardiomyocyte-specific promoter^23^.

In this work, we similarly combine the temporal control over gene expression offered by a tet-ON tetracycline-inducible system with the power of Cre recombination under the control of a cell-type specific promoter to generate cardiomyocyte-specific, inducible, reprogrammable mice – herein also referred to as Myh6-Cre^+^Col1a1^OKSM^ mice – and show the ability of adult mammalian cardiomyocytes to undergo reprogramming. We demonstrate that OKSM expression is induced exclusively in cardiomyocytes by oral administration of doxycycline in this model, and that these cells undergo complete reprogramming to *bona fide* pluripotency in situ when doxycycline is administered for eighteen days in a row. Moreover, we demonstrate that partial reprogramming without tumorigenesis is also feasible by inducing short cycles of ON and OFF OKSM expression, and that this strategy results in cardiomyocyte epigenetic rejuvenation which correlates with the amelioration of a cardiac failure phenotype at the functional level.

## RESULTS

### A cardiomyocyte-specific, inducible, reprogrammable mouse

To investigate if adult mouse cardiomyocytes can undergo OKSM-induced reprogramming *in vivo*, we generated triple mutant mice in which the expression of the four reprogramming factors is induced exclusively in cardiomyocytes and in response to a chemical stimulus (**Fig. 1a**). In brief, cardiomyocyte-specific expression of Cre recombinase, driven by the cardiomyocyte-specific alpha myosin-heavy chain *Myh6* promoter^24^, was combined with a tet-ON doxycycline-inducible system to control the expression of OKSM^25^ and conditional expression of the reverse tetracycline-controlled transactivator (rtTA)^26^. rtTA and GFP are preceded by a STOP sequence flanked by loxP sites – thus excised only in cardiomyocytes, by Cre – and, in the presence of doxycycline, rtTA induces OKSM expression. Fluorescence activated cell sorting (FACS) of different cardiac cell types (cardiomyocytes, cardiac fibroblasts, endothelial cells and leukocytes) demonstrated that GFP expression – and thus Cre recombination – is restricted to cardiomyocytes of Myh6-Cre^+^ mice (**Fig. 1b-d**). We found a very small percentage (0.46%) of endothelial cells that fluoresced in the GFP gate; however, those are likely just cells with a higher level of autofluorescence since a similar percentage (0.47%) was found among the CD31^+^ cells in control Myh6-Cre^−^ mice (**Supplementary Fig. 1**). To obtain a further proof of the cardiomyocyte specificity of the system, 1 mg/ml doxycycline was administered in the drinking water of Myh6-Cre^+^Col1a1^OKSM^ mice for a week and cardiomyocytes, cardiac endothelial cells and the rest of non-myocyte cells – containing likely an enrichment of cardiac fibroblasts and some immune cells – were isolated with a combination of centrifugation and magnetic cell sorting (**Fig. 1e**). Real-time, reverse-transcription, quantitative PCR (RT-qPCR) analysis demonstrated that *Oct3/4* and *GFP* were significantly expressed in cardiomyocytes only (**Fig. 1f-g**). To confirm the purity of the different cell types isolated, the expression levels of cell-specific markers *Myhl2*, *Ddr2* and *Vwf* – a cardiomyocyte, cardiac fibroblast and endothelial cell specific marker, respectively – were measured and found to be significantly upregulated in the respective cell fractions (**Supplementary Fig. 2**). Analysis of other organs, including the liver (**Supplementary Fig. 3b**), spleen (**Supplementary Fig. 3c**) and kidney (**Supplementary Fig. 3d**) confirmed the lack of significant off-target expression of the GFP and OKSM cassettes aside from very low levels of GFP in the liver. However, this was not considered to compromise the specificity of our model since it was not accompanied by the expression of *Oct3/4* or significant upregulation of *Sox2*, which would be indicative of efficient recombination.

**Fig. 1.**
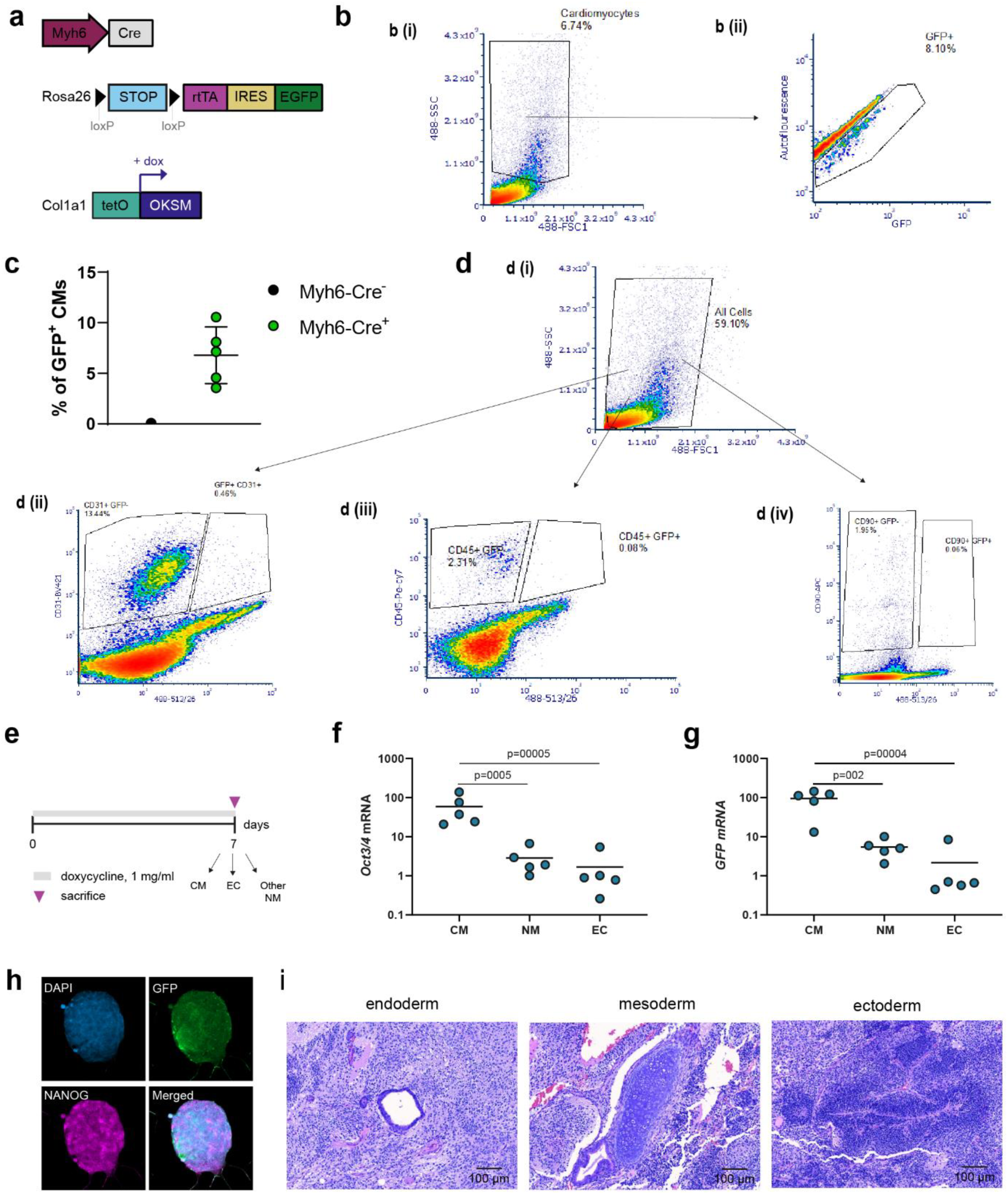
Generation and characterization of cardiomyocyte-specific, inducible, reprogrammable mice. **(a)** Schematic representation of the gene circuits utilized to create cardiomyocyte-specific, inducible, reprogrammable mice. **(b)** Representative FACS sorting plots of GFP^+^ cardiomyocytes from Myh6-Cre^+^Col1a1^OKSM^ mice (n=5). (**bi**) Shows sorting of cardiomyocytes based on their larger size and (**bii**) shows the population of GFP^+^ cells within the cardiomyocyte fraction. **(c)** Percentage of GFP^+^ cardiomyocytes in Myh6-Cre^−^Col1a1^WT^ mice (n=2) and Myh6-Cre^+^Col1a1^OKSM^ mice (n=5) obtained from FACS studies. **(d)** Sorting of GFP^+^ cells from cardiac cell populations other than cardiomyocytes. (**di**) Shows gating on all cardiac cells, (**dii**) CD31^+^ endothelial cells, (**diii**) CD45^+^ leukocytes and (**div**) CD90^+^ fibroblasts. **(e)** Schematic representation of the experimental design to analyze gene expression in different cardiac cell types after doxycycline administration. mRNA levels of the reprogramming factor *Oct3/4* **(f)** and *GFP* **(g)** in different cardiac cell types after 7 days of doxycycline administration. CM: cardiomyocyte, NM: non-myocyte, EC: endothelial cell. Gene expression levels were normalized to those in cardiomyocytes. Individual data points represent the fold change (2^^−ΔΔCt^) of each replicate (n=5 mice/group). Statistical analysis was performed by one-way ANOVA and Tukey’s post-hoc test. **(h)** Immunocytochemistry of CM-iPSC colonies. Images were taken at 20X magnification, scale bar = 50 μm. **(i)** Endoderm, mesoderm and ectoderm representation in H&E stained sections of teratomas obtained upon inoculation of CM-iPSC colonies in immunodeficient mice. Images were taken at 10X, scale bar represents 100 μm.

We also confirmed that cardiomyocytes isolated from the hearts of adult Myh6-Cre^+^Col1a1^OKSM^ mice can be reprogrammed into induced pluripotent stem cells (iPSCs) in culture (**Fig. 1h-i** and **Supplementary Fig. 4**). This is, to our knowledge, the first report of adult cardiomyocyte reprogramming into iPSCs *in vitro*, which was unsuccessful in a previous attempt^23^. Cardiomyocytes were isolated from other cardiac cells following a previously described Langendorff-free method and several rounds of gravity sedimentation^27^. Colonies started to form after two weeks in culture supplemented with doxycycline and leukemia inhibitory factor (LIF), in cardiomyocyte cultures only, and stained positively for the pluripotency marker SSEA1 shortly after (**Supplementary Fig. 4a)**. After doxycycline withdrawal, we confirmed characteristic dome shaped morphology (**Supplementary Fig. 4a)**, co-expression of GFP with reprogramming and pluripotency markers (**Fig. 1i** and **Fig.S4b)** and the capability of cardiomyocyte-derived iPSCs (CM-iPSCs) to form teratomas with trilineage contribution and positive GFP staining when subcutaneously inoculated into immunodeficient mice (**Fig. 1h** and **Supplementary Fig. 4c-d**). Overall, these results show that our mouse model enables cardiomyocyte-specific induction of OKSM and that adult cardiomyocytes can be reprogrammed under specific culture conditions.

### Sustained OKSM expression establishes a genetic program characteristic of pluripotency and induces cardiomyocyte de-differentiation *in situ*

The drinking water of reprogrammable (Myh6-Cre^+^Col1a1^OKSM^) or control (Myh6-Cre^+^Col1a1^WT^) mice was next supplemented with 1 mg/ml doxycycline for 7, 12 and 18 days, and changes in gene expression in the ventricular myocardium were analyzed (**Fig. 2a**). Increased expression of OKSM factors was detected by day 7 in reprogrammable mice treated with doxycycline, and further increased on day 12 to reach a plateau (**Fig. 2b-c**). On day 12, a slight, albeit not statistically significant, elevation was found in the mRNA levels of pluripotency-related genes, including *Nanog*, the endogenous form of *Oct4* (*endoOct4*) and *Gdf3* in the reprogrammable group (**Fig. 2d**). These genes, which are typically silenced in adult differentiated cells but re-expressed from the maturation phase of induced pluripotent stem cell (iPSC) reprogramming onwards ^28^, were significantly and very highly upregulated on day 18 (>1000 fold, on average, for *Nanog* and >100 fold, on average, for *endoOct4* and *Gdf3*). We confirmed that these changes were only observed in reprogrammable mice (Myh6-Cre^+^Col1a1^OKSM^) administered with doxycycline and not in mutants lacking Cre recombination (Myh6-Cre^−^Col1a1^OKSM^) or the OKSM cassette (Myh6-Cre^+^Col1a1^WT^), or in reprogrammable mice that were administered unadulterated, normal drinking water (**Supplementary Fig. 5**). Raising the dose of doxycycline to 2 mg/ml did not trigger higher mRNA expression levels of the OKSM reprogramming factors (on day 18) or of the endogenous pluripotency genes investigated, at any of the time points analyzed (**Supplementary Fig. 6**). Therefore, the 1 mg/ml doxycycline dose was maintained for the rest of the studies.

**Fig. 2.**
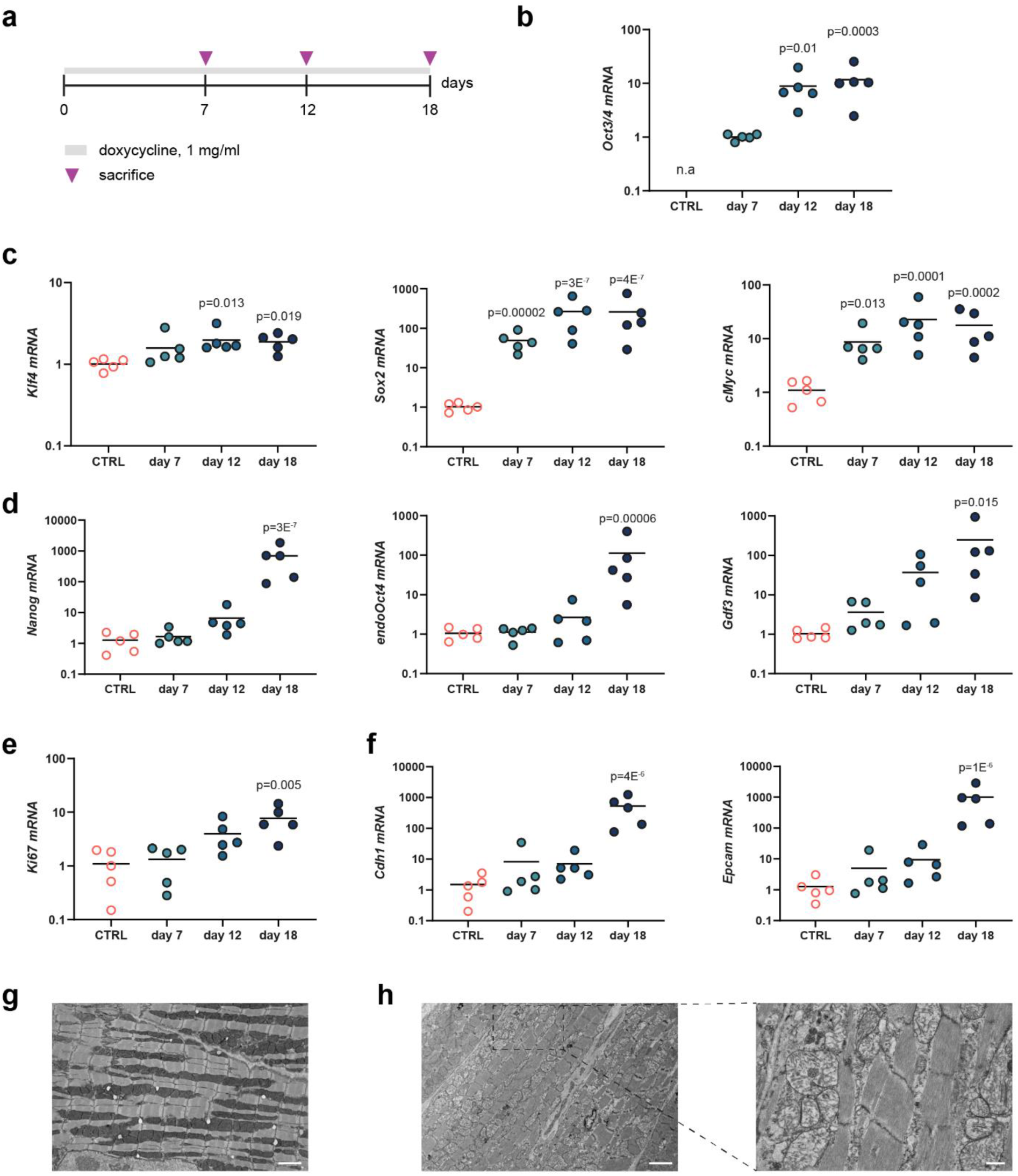
Eighteen days of doxycycline-induced OKSM establish a gene expression program characteristic of pluripotent reprogramming in the myocardium and induces cardiomyocyte de-differentiation. **(a)** Schematic representation of experimental design. **(b)** mRNA levels of *Oct3/4* reprogramming factor. **(c)** mRNA levels of other reprogrammable factors, *Klf4*, *Sox2* and *cMyc*. **(d)** Gene expression levels of endogenous pluripotency markers. **(e)** Gene expression levels of *Ki67*. **(f)** mRNA levels of MET transition markers. *Oct3/4* levels were normalized to those in Myh6-Cre^+^Col1a1^OKSM^ mice (7 days of doxycycline). The expression of all other genes investigated was normalized to that of control Myh6-Cre^+^Col1a1^WT^ mice. Individual data points represent the fold change (2^^−ΔΔCt^) of each replicate (n=5 mice/group). Statistical analysis was performed by Welch ANOVA and Games-Howell post-hoc test (*Gdf3* data) or one-way ANOVA and Tukey’s post-hoc test (all other genes). P values refer to statistical significance in relevance to the control group. **(g)** Representative TEM micrograph of cardiac ventricle tissue from reprogrammable mice that received normal drinking water, scale bar denotes 2 µm. **(h)** Representative TEM micrographs of cardiac ventricle tissue from reprogrammable mice administered with 1 mg/ml doxycycline in the drinking water for 18 days. Scale bars represent 2 µm (left) and 500 nm (closeup, right). The figure shows representative images from 5 random fields of view (FOV) and n=2 mice/group.

We then investigated other signs of reprogramming at the gene expression level. A significant upregulation of *Ki67*, a marker of cell cycle activity, was found after 18 days of doxycycline-induced OKSM reprogramming (**Fig. 2e**), in agreement with the increased proliferation rate that is observed during cell reprogramming ^4^. There was also a significant increase of *Cdh1* and *Epcam* mRNA levels – markers of the mesenchymal-to-epithelial transition which is also characteristic of pluripotent reprogramming – at the same time point (**Fig. 2f**). Lastly, the levels of ten-eleven translocation methyl cytosine dioxygenases (TET) enzymes, known to catalyze DNA demethylation during cell reprogramming, were also significantly upregulated in reprogrammable mice during the course of the study (**Supplementary Fig. 7**). Lu et al have recently shown that active demethylation catalyzed by these enzymes is indeed necessary for OKS expression to translate into regenerative effects ^6^.

The architecture and morphology of reprogrammed cardiac tissues was also analyzed by transmission electron microscopy (TEM) for signs of reprogramming. The ventricular myocardium of reprogrammable mice that received drinking water without doxycycline showed the typical structure of aligned cardiac muscle fibers with organized sarcomeres and clearly distinguishable A and I bands and Z lines (**Fig. 2g**, **Supplementary Fig. 8b**). However, ventricular tissues from reprogrammable mice that were offered doxycycline water for 18 days contained large areas filled with de-differentiated cardiomyocytes, as evidenced by the loss of fiber organization and sarcomere assembly (**Fig. 2h**). Sarcomere disassembly was also observed, albeit with less frequency, after 12 days of doxycycline administration (**Supplementary Fig. 8c**). On days 3 and 7 after the start of OKSM induction, no differences were observed, as compared to mice that did not receive doxycycline (**Supplementary Fig. 8c**).

### Eighteen days of doxycycline-induced OKSM expression induces stable cardiomyocyte reprogramming to pluripotency

Moved by the high expression pf pluripotency markers detected in the tissue (**Fig. 2d**), we next set to investigate if 18 days of doxycycline-induced OKSM expression were sufficient to trigger the conversion of cardiomyocytes into stable (i.e., independent from exogenous OKSM) and functionally pluripotent stem cells in situ. Previous studies in mouse liver and skeletal muscle have shown that *in vivo* reprogrammed cells can exhibit some molecular hallmarks of pluripotency transiently without committing to pluripotency ^1, 3, 29^. We administered doxycycline-supplemented water to reprogrammable (Myh6-Cre^+^Col1a1^OKSM^, n=8) and control (Myh6-Cre^−^Col1a1^OKSM^, n=10; Myh6-Cre^+^Col1a1^WT^, n=14) mice for 18 days, followed by withdrawal of the drug (**Fig. 3a**). No significant changes in body weight were observed for the first 30 days of the study (**Fig. 3b**), after which reprogrammable mice had to be euthanized for humane reasons (mainly showing difficulty in breathing and lethargy). The entire group succumbed within 58 days of the start of doxycycline treatment (**Fig. 3d**). Upon necropsy investigation, 75% (6/8) of Myh6-Cre^+^Col1a1^OKSM^ mice showed clear tumor masses growing out of the heart (**Fig. 3c, e**). One mouse in that group was found dead on day 18 and another one had to be euthanized on day 58 due to a spontaneous tumor growing on the head, but histological investigation did not reveal any signs of dysplasia or teratoma formation in the heart in either of those cases (data not shown). On the contrary, mice from the control Myh6-Cre^−^Col1a1^OKSM^ group survived for the expected lifespan of the C57BL/6 background (**Fig. 3d**) and did not show any evidence of cardiac tumorigenesis upon necropsy. The reduced lifespan of Myh6-Cre^+^Col1a1^WT^ mice (**Fig. 3d**) is likely linked to the continued expression of Cre recombinase and GFP in cardiomyocytes, which has been previously described as cardiotoxic and a limitation of transgenic models ^30–32^ but, as in the Myh6-Cre^−^Col1a1^OKSM^ group, no cardiac teratomas were observed at the time of death. In agreement with previous reports, we confirmed that Myh6-Cre^+^Col1a1^WT^ mice develop age-dependent dilated cardiomyopathy with significant decline of systolic heart function from 8 months of age (**Supplementary Fig. 9**).

**Fig. 3.**
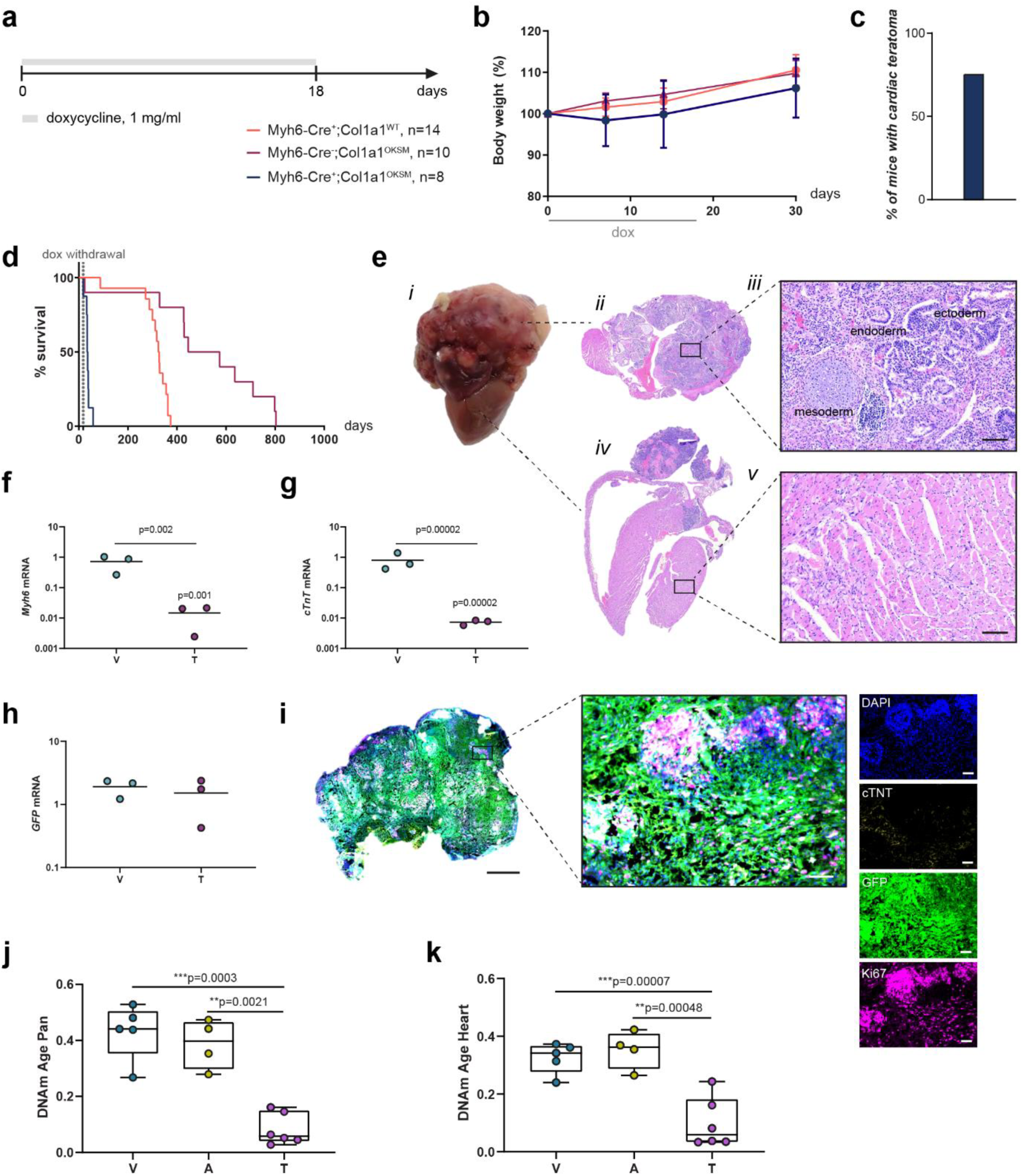
Doxycycline-induced expression of OKSM for 18 days reprograms cardiomyocytes to functionally pluripotent cells that generate teratomas. **(a)** Schematic representation of experimental design. **(b)** Changes in body weight upon administration of 1 mg/ml doxycycline in the drinking water of reprogrammable (Myh6^+^Col1a1^OKSM^) and control (Myh6^+^Col1a1^WT^; Myh6^−^ Col1a1^OKSM^) mice. **(c)** Incidence of teratoma formation upon doxycycline treatment. Teratomas were only found in reprogrammable mice. **(d)** Survival of reprogrammable and control mice after 18 days on doxycycline treatment. **(e)** Representative teratoma in the heart of a reprogrammable mouse (i): whole tissue section (ii) and close-up at 10X magnification (iii) of teratoma H&E staining; whole tissue section (iv) and close-up at 10X magnification (v) of the four chambers of the heart. Scale bars represent 100 µm. Gene expression levels of *Myh6* (f), *cTnT* (g), and *GFP* (h) in cardiac teratomas and ventricular myocardium. Gene expression data are normalized to that of Myh6^+^Col1a1^WT^ cardiac tissue exposed to the same doxycycline regime. Individual data points represent the fold change (2^^−ΔΔCt^) of each replicate (n=5 mice/group). Statistical analysis was performed by one-way ANOVA and Tukey’s post-hoc test. P values refer to statistical significance in relevance to the control group or between ventricle and teratoma (*Myh6* and *cTnT*, denoted with a line between the two groups). **(i)** Triple immunofluorescence for cTNT, GFP and Ki67 in cardiac teratomas. Scale bar represents 1 mm (whole section) and 50 µm (close-up at 10X magnification). DNA methylation age of bulk ventricle (V), atrium (A) and teratoma (T) tissue isolated from Myh6^+^Col1a1^OKSM^ mice calculated with the application of a pan-tissue **(j)** and heart-specific **(k)** epigenetic clock.

Histological investigation of reprogrammed hearts confirmed trilineage contribution (i.e., teratoma identity) in tumor masses sprouting from the heart (**Fig. 3e, *ii-iii***) and other areas of dysplasia were found in atrial and ventricular tissues (**Fig. 3e, *iv-v***). We then studied gene and protein expression patterns in the teratomas to lineage trace their origin. *Myh6* (**Fig. 3f**) and *cTnT* (**Fig. 3g**) mRNA levels were significantly downregulated in teratomas compared to their respective ventricular tissue but GFP expression levels were maintained (**Fig. 3h**), suggesting that cells within teratomas originated from cardiomyocytes that lost their identity upon reprogramming. These observations were confirmed by immunohistochemistry (**Fig. 3i**). Teratomas stained strongly for GFP and showed clusters of proliferative Ki67^+^ cells, while the expression of cTNT was extremely low and not uniformly detected. In addition, the expression of OKSM factors (**Supplementary Fig. 10a**), endogenous pluripotency markers (**Supplementary Fig. 10b**) and a significant upregulation of *Ki67* (**Supplementary Fig. 10c**) were confirmed in teratomas and ventricular myocardium of reprogrammed hearts.

Ageing is accompanied by changes to DNA methylation at specific CpG sites of the mammalian genome. Likewise, the biological age of tissues grows with age, whereas the predicted age of iPSCs is close to zero^33, 34^. We used epigenetic clocks – machine learning algorithms trained on DNA methylation data^33, 35, 36^ – to quantify the inferred biological age of the atria (A), ventricles (V) and teratomas (T) of reprogrammable mice. Both a pan-tissue (**Fig. 3j**) and heart-specific (**Fig. 3k**) clocks^37–39^ showed that the DNA methylation age of cardiac teratomas significantly decreased to values close to zero, whereas we observed no difference between ventricle and atrium.

We also performed a classic teratoma assay, a gold-standard assay of functional pluripotency^40^, to confirm the cardiomyocyte origin and differentiation potential of *in vivo* reprogrammed cells. Reprogrammable (Myh6-Cre^+^Col1a1^OKSM^, n=7) or control mice (Myh6-Cre^+^Col1a1^WT^, n=4) were administered doxycycline-supplemented water for 18 days, after which they were sacrificed, their atrial and ventricular chambers digested separately into single cell suspensions and injected subcutaneously (s.c.) in opposite flanks of NU/J immunodeficient mice (**Supplementary Fig. 11a**). After 4 weeks, all cell suspensions obtained from atrial reprogrammed tissue (7/7) and 29% of those from the ventricles (2/7) had generated GFP+ teratomas (**Supplementary Fig. 11b-g**). No teratomas resulted from the inoculation of cardiac cell suspensions obtained from control, non-reprogrammable mice (**Supplementary Fig. 11b**). Given the higher incidence of atrial teratomas, we analyzed changes in gene expression and morphology of this tissue upon the administration of doxycycline (**Supplementary Fig. 12** and **Supplementary Fig. 13**). The expression of reprogramming and endogenous pluripotency genes followed a very similar trend as seen in the ventricles (**Supplementary Fig. 12b-c**). Changes in sarcomere structure were also evident from day 12 after the start of the doxycycline treatment but, overall, there were more areas showing clear cardiac muscle fiber de-differentiation in the atria than in the ventricles (**Supplementary Fig. 13**). These results confirm that cardiomyocyte reprogramming takes place in all heart chambers in these mice, with likely a higher efficiency of reprogramming in the atria. This is not surprising, as *Myh6* is expressed in both atrial and ventricular tissue in mice, albeit with varying levels throughout development^41^.

Overall, our results show that adult cardiomyocytes can be reprogrammed *in vivo* and that they commit to pluripotency after 18 days of continuous OKSM expression, even when the forced upregulation of reprogrammed factors is interrupted, and while they are maintained in the native microenvironment of the adult mouse myocardium.

### Cyclic upregulation of OKSM prevents re-acquisition of pluripotency and teratoma formation

The induction of cardiomyocyte complete reprogramming to pluripotency *in vivo* provides important insights into the plasticity of post-mitotic cells, however, it lacks clinical relevance due to the generation of teratomas. This motivated us to identify suitable OKSM upregulation regimes that would result in partial reprogramming without reacquisition of pluripotency and risk of tumorigenesis. We tested a 2-day-doxycycline-ON/5-day-doxycycline-OFF protocol (**Fig. 4**), herein referred to as 2ON/5OFF, that triggered partial reprogramming and erasure of aging hallmarks in a previous study with a different reprogrammable mouse model ^5^. We first administered six cycles of this regime and followed survival and appearance of teratomas upon withdrawal of the drug (**Fig. 4a**). Reprogrammable mice subjected to this protocol survived for their expected lifespan (**Figs. 4b-c**) and none of them showed macroscopic signs of teratoma formation in the heart at the time of death. We then investigated gene expression and tissue architecture at the end of the ON phase of the last (6^th^) cycle (**Fig. 4d**). RT-qPCR data confirmed that OKSM reprogramming factors were upregulated in the ventricles of doxycycline-administered reprogrammable mice at fold changes similar to those observed after 18 days of continued drug administration (**Figs. 4e-f**). However, the levels of the endogenous pluripotency genes *Nanog*, *Gdf3* and *endoOct4* remained unchanged compared to non-reprogrammable controls (**Fig. 4g**). Similarly, no changes were observed in the expression of MET markers (**Fig. 4h**) and of *Ki67* (**Fig. 4i**). At the histological level, no apparent de-differentiation, dysplasia or overall changes to fiber morphology and sarcomere organization were detected in the ventricles (**Fig. 4j**) or atria (**Fig. 4k**) of reprogrammable mice compared to controls. These data confirm that cyclic doxycycline administration induces significant OKSM expression in the heart without pluripotent conversion.

**Fig. 4.**
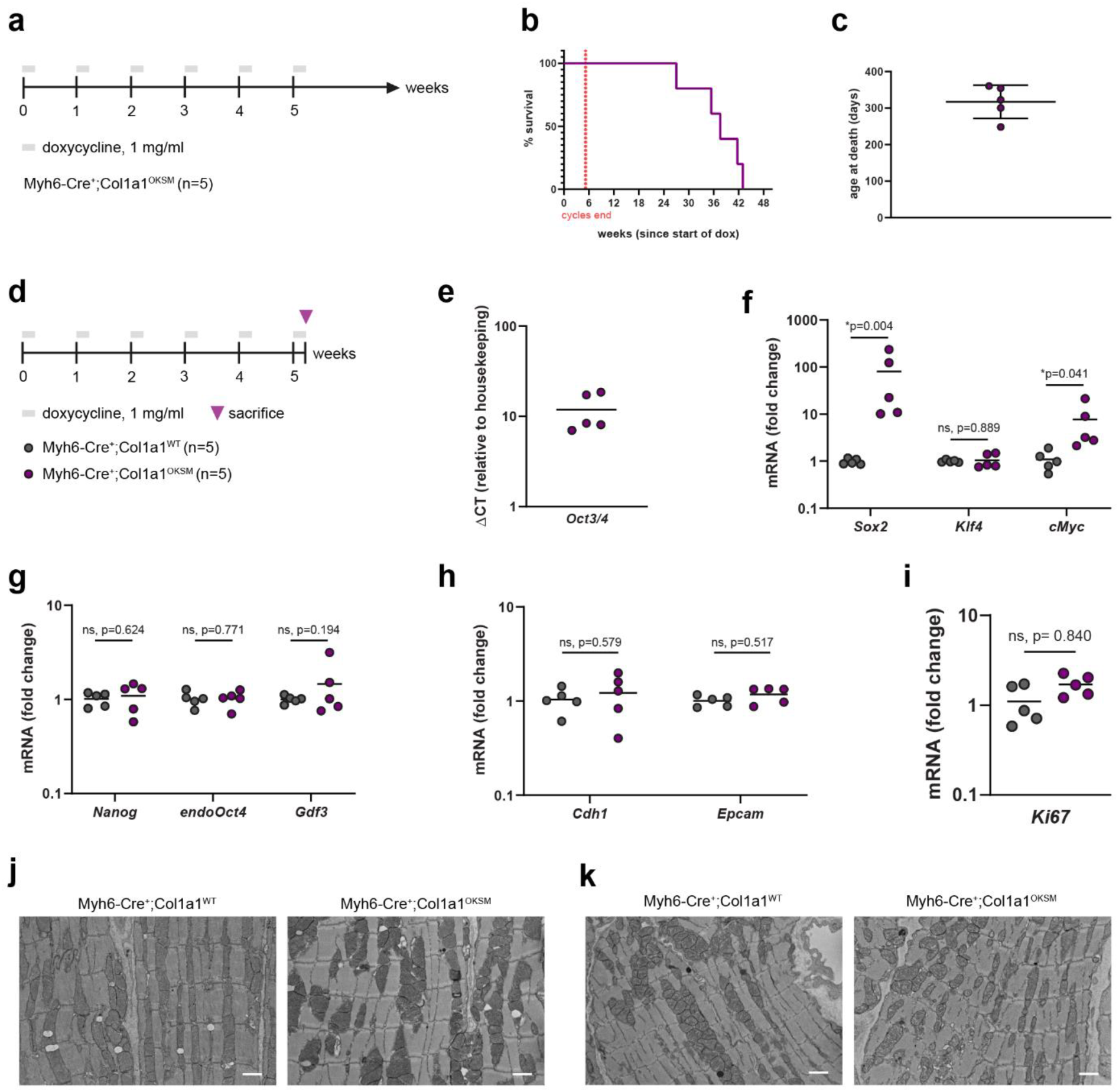
Cyclic OKSM induction (2-day-ON/5-day-OFF) prevents pluripotency re-acquisition and teratoma formation. **(a)** Schematic representation of experimental design, survival study. **(b)** Survival of reprogrammable mice (Myh6-Cre^+^Col1a1^OKSM^) administered with 6 cycles of 2ON/5OFF doxycycline, n=5. **(c)** Age at the time of death of reprogrammable mice (Myh6^+^Col1a1^OKSM^) administered with 6 cycles of 2ON/5OFF doxycycline. **(d)** Schematic representation of experimental design, gene expression and histology study. Gene expression levels of reprogramming factors *Oct3/4* **(e)**, *Klf4, Sox2 and cMyc* **(f)**, endogenous pluripotency markers **(g)**, markers of MET **(h)** and *Ki67* **(i)**. Gene expression data are normalized to that of Myh6-Cre^+^Col1a1^WT^ cardiac tissue exposed to the same doxycycline regime, except for *Oct3/4* mRNA levels. The latter were normalized to the geometric mean of two reference genes (*Mapk1* and *Rps13*). Individual data points represent the fold change (2^^−ΔΔCt^) of each replicate (n=5 mice/group). Statistical analysis was performed by one-way ANOVA and Tukey’s post-hoc test. TEM micrographs of ventricular **(j)** and atrial **(k)** tissues of Myh6^+^Col1a1^WT^ and Myh6^+^Col1a1^OKSM^ mice treated with 6 cycles of 2ON/5OFF doxycycline. Scale bars represent 500 nm.

### Cyclic OKSM induction in 9-month-old mice triggers cardiomyocyte epigenetic rejuvenation and correlates with the amelioration of cardiac failure

We next decided to investigate the effects of the 2ON/5OFF regime in reprogrammable mice at 9 months of age, when they have already developed functional signs of cardiac failure (**Supplementary Fig. 9**). All our previous reprogramming experiments were performed on 8 to 12-week-old mice. 9-month-old reprogrammable mice (Myh6-Cre^+^Col1a1^OKSM^) and control littermates (Myh6-Cre^−^Col1a1^WT^) were administered 1 mg/ml doxycycline following the 2ON/5FF regime (6 cycles) or provided with sucrose water only as controls (**Fig. 5a**). Systolic heart function was monitored just before and at the end of the study. Mice were sacrificed at the end of the ON phase of the 6^th^ cycle – when they were 10.5 months old – and cardiomyocytes and non-myocytes were collected separately for analysis to avoid small changes expected in cardiomyocytes only being masked in a bulk tissue analysis. For these studies, young (3-month-old) counterparts were also included as controls. We first used RT-qPCR to confirm that OKSM factors were upregulated in cardiomyocytes from these older reprogrammable mice (**Fig. 5b**) while endogenous pluripotency genes remained at baseline levels (**Fig. 5c**), in agreement with our data in younger mice (**Figs. 4e-g**). We then applied a heart-specific epigenetic clock to investigate changes in DNA methylation age of reprogrammed and control cardiomyocytes (**Fig. 5d**). Importantly, the administration of doxycycline did not affect DNA methylation age by itself, as evidenced by the absence of significant differences between treated and untreated Myh6-Cre^−^Col1a1^WT^ cardiomyocytes. On the contrary, we observed a significant decrease of epigenetic age in doxycycline-treated cardiomyocytes from reprogrammable mice, compared to their untreated counterparts, which suggests that cyclic OKSM can induce significant cardiomyocyte rejuvenation at the epigenetic level. Changes in epigenetic age were not detected in the non-myocytes of Myh6-Cre^+^Col1a1^OKSM^ mice (**Fig. 5e**), highlighting once again the cardiomyocyte specificity of our model. These findings are robust, as similar trends were confirmed with two different pan-tissue clocks (**Supplementary Fig. 14**).

**Fig. 5.**
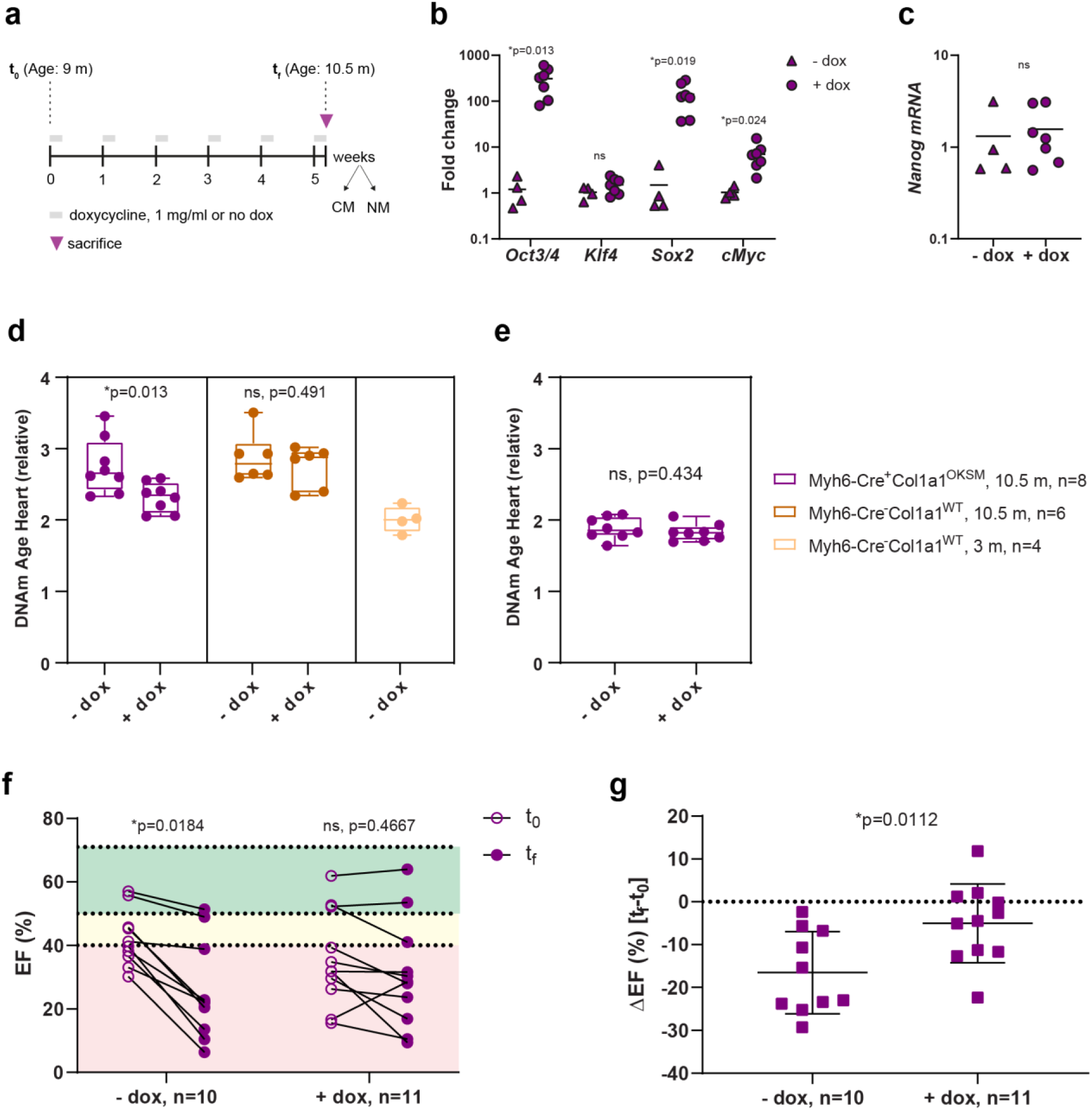
Cyclic OKSM induction (2-day-ON/5-day-OFF) reduces cardiomyocyte epigenetic age and halts the decline of cardiac function in aged Myh6^+^Col1a1^OKSM^ mice. **(a)** Schematic representation of experimental design. Doxycycline treatment (2ON/5OFF, 6 cycles) was initiated at 9 months of age and systolic heart function (ejection function, EF) was assessed via echocardiography at the start (t_0_) and end of the study (t_f_). Cardiomyocytes (CM) and non-myocytes (NM) were isolated at the end of the ON phase of the 6^th^ cycle. mRNA levels of **(b)** OKSM factors and (c) *Nanog*, measured by RT-qPCR. Heart-specific epigenetic clock applied to **(d)** cardiomyocyte and **(e)** non-myocyte DNA methylation profiles. Ejection fraction (EF) values at t_0_ and t_f_ are shown in **(f)** and expressed as the ΔEF (t_f_-t_0_) in **(g)**. P values were calculated by independent samples t tests and represent statistically significant differences between the t_f_ and t_0_ (in f) and between the ΔEF in treated and untreated groups (in g). _*_p<0.05, ns= not significant.

Lastly, we investigated changes in systolic heart function by high-frequency ultrasound imaging (**Fig. 5a**). Ejection fraction (EF) values remained unchanged in the Myh6-Cre^−^Col1a1^WT^ group regardless of the administration of doxycycline (**Supplementary Fig. 15**). Moreover, these mice presented physiological EF values, as expected. On the contrary, Myh6-Cre^+^Col1a1^OKSM^ mice presented EF values characteristic of severe cardiac failure from the beginning of the study (t_0_, 9 months of age, **Fig. 5f**). At the end of the study (t_f_, 10.5 months of age), EF had dropped significantly in untreated Myh6-Cre^+^Col1a1^OKSM^ mice while it remained stable in the doxycycline-treated group (**Figs. 5f-g**). This suggests that while partial cardiomyocyte reprogramming and epigenetic rejuvenation may not be able to improve cardiac function in this severe condition, it may at least be able to contribute to stopping or slowing further functional decline. Overall, our data demonstrate that cyclic OKSM induction in aged cardiomyocytes results in a significant reduction of their epigenetic age, without pluripotent conversion or tumorigenesis, and this correlates positively with the amelioration of a severe cardiac failure phenotype at the functional level. Further studies to investigate the relationship between epigenetic rejuvenation and this functional improvement of the cardiac muscle are warranted.

## DISCUSSION

We demonstrate here that Cre recombination under the control of a cell type-specific promoter and conditional, doxycycline-inducible regulation of gene expression can be combined to achieve cardiomyocyte-specific expression of OKSM and reprogramming. Using this strategy, we have confirmed that adult mouse cardiomyocytes can be fully reprogrammed to the pluripotent state *in vivo*. This possibility had remained elusive in earlier studies – likely due to a combination of low OKSM expression, poor temporal control over the latter, and use of viral vectors with promiscuous tropism to induce OKSM^17^ – until a recent report by Chen et al^23^ that used a reprogrammable mouse model similar to the one described here. The ability to reverse the fully differentiated state of adult mammalian cardiomyocytes to *bona fide* pluripotency is an important observation, since there is typically an inverse relationship between the differentiation status of a cell and its capacity to undergo reprogramming^19–21^ and reports of post-mitotic cells undergoing reprogramming are scarce, mainly limited to transdifferentiation events^42, 43^ or require p53 suppression to force cell proliferation^44^.

Our studies also confirm that pluripotency and tumorigenesis can be avoided by limiting the duration of OKSM expression. We show here that cardiomyocytes from mice exposed to short cycles of OKSM upregulation retain the silencing of the pluripotency network characteristic of differentiated cells, but their DNA methylation profile reflects that of a chronologically younger age. These findings agree with other reports that short expression of OKSM (in cycles or as a single event) erase several hallmarks of cellular aging^5, 6, 12, 13, 45^. Ours is, to our knowledge, the first report of OKSM-triggered cardiomyocyte epigenetic rejuvenation. In addition, our observations suggest a positive correlation between this effect and the stabilization of a severe cardiac condition at the functional level. However, this observation should be validated in more physiologically relevant models of cardiac disease, and the mechanism or mechanisms by which partially reprogrammed cardiomyocytes may be contributing remain to be explained.

Importantly, our findings and those by Chen and colleagues^23^ suggest a new avenue for cardiac regeneration through cardiomyocyte reprogramming although with significant differences likely enabled by the step-wise character of the reprogramming process. Chen et al used a different cardiomyocyte-specific reprogrammable mouse (with Cre under the control of the *Myl2* promoter) and short (6-day-long) rather than cyclic administration of doxycycline and observed cardiomyocyte proliferation and neonatal-like gene expression associated with an improvement of regeneration after myocardial infarction^23^. Their approach adds to other gene therapies also proposed to stimulate endogenous cardiomyocyte cell division, including those leveraging cell cycle regulators such as Cyclin D2^46^, metabolic regulators such as ERRB2 ^47^ or non-coding RNAs that regulate cardiomyocyte regeneration in other species^48^. In contrast, under our experimental protocol we did not find evidence of cell cycle re-entry. Our data shows, however, epigenetic rejuvenation of cardiomyocytes and positive correlation with the amelioration of age-dependent heart failure, which suggests that the therapeutic potential of partial cardiomyocyte reprogramming may not be limited to the repopulation of lost cardiomyocytes. Instead, it may also offer opportunities to treat and prevent ailments of the aging heart that do not necessarily present with extensive cardiomyocyte death. Indeed, others have shown (in other tissues) that shorter OKSM expression or the absence of *cMyc* limit reprogramming to the resetting of ageing signatures without proliferation, and this rejuvenation has shown to increase overall fitness and improve the outcomes of subsequent tissue damage^5, 6^. Reprogramming without cell cycle re-entry may also be a safer approach as long as new cells are not needed in the damaged tissue, since it would eliminate the risk of uncontrolled or excessive proliferation. Overall, the fact that beneficial effects linked to cardiomyocyte reprogramming have been found independently and through different mouse models and reprogramming protocols is extremely encouraging.

Cell type-specific reprogrammable mouse models may enable higher mechanistic depth of studies, for example, to elucidate key cell types required to be reprogrammed for the successful regeneration of a specific tissue, or to investigate the crosstalk between reprogrammed and non-reprogrammed cells, but examples of these models are still scarce in the scientific literature. Besides the work of Chen et al and this study, there are only two other reports of reprogrammable models with cell type specificity^9, 10^. Interestingly, Wang et al uncovered a paracrine effect of partially reprogrammed myofibers on the skeletal muscle stem cell niche as the direct mediator of muscle regeneration, which could not have been elucidated using traditional reprogrammable mice with ubiquitous OKSM expression. Future research in this field will also likely focus on the development of clinically translatable alternative approaches. Doxycycline-inducible adeno-associated viral vectors have shown promise towards this goal thanks to tissue-specific tropism and control over the duration of gene expression^6^ but they still do not offer tropism with cell type resolution unless combined with Cre transgenic lines. In parallel, extensive research on small molecule drugs to replace OKSM is underway^49^.

In sum, our findings support the ability of adult cardiomyocytes to be reprogrammed to pluripotency or to epigenetically rejuvenated states, and more broadly the potential of *in vivo* cell reprogramming for regenerative medicine and a platform with high control over OKSM expression to enable further mechanistic studies.

## EXPERIMENTAL PROCEDURES

### Mice

All animal procedures were performed in accordance with the Harvard University Faculty of Arts and Sciences (FAS) Institutional Animal Care and Use Committee (IACUC) guidelines. B6.FVB-Tg(Myh6-cre)2182Mds/J mice express constitutively active Cre recombinase enzyme under the control of the Myh6 promoter through a transgene insertion^24^, and were obtained from the Jackson Laboratory (Cat #011038). B6.Cg-*Gt(ROSA)26Sor^tm1(rtTA,EGFP)Nagy^*/J mice contain a floxed (flanking loxP) Pgk-neo-pA cassette and a rtTA-IRES-EGFP-pA cassette inserted into intron 1 of the Rosa26 locus^26^, and were purchased from the Jackson Laboratory (Cat #005670). B6;129S4-*Col1a1^tm1(tetO–Pou5f1,–Klf4,–Sox2,–Myc)Hoch^*/J mice have a targeted mutation in the *Col1a1* gene. The OKSM cassette consisting of four mouse reprogramming genes, *Oct3/4*, *Klf4*, *Sox2*, and *cMyc*, is expressed under the control of the tet-responsive element (*tetO*) with CMV minimal enhancer-less promoter^25^. These mice were a kind gift from Prof Konrad Hochendlinger (Massachusetts General Hospital and Harvard University). The three strains above were bred *in house* to generate cardiomyocyte-specific, inducible reprogrammable mice. The rtTA-IRES-EGFP-pA was kept in homozygosis while the Cre recombinase and OKSM modifications were maintained as heterozygotes following advice from donating investigators for optimal breeding. For simplicity, triple transgenic reprogrammable mice are named Myh6-Cre^+^Col1a1^OKSM^ or cardiomyocyte-specific reprogrammable mice hereinafter. Myh6-Cre^+^Col1a1^WT,^ Myh6-Cre^−^ Col1a1^OKSM^ and Myh6-Cre^−^Col1a1^WT^ mice were included as *controls* throughout the study, as specified for each experiment. Mice of 8-12 weeks of age and of both sexes were used for all reprogramming studies, unless otherwise specified. Male mice of 3 or 9 months of age were used in the rejuvenation studies in **Fig. 5** and **Supplementary Figs. Fig.14-15**, and male mice of 6, 8, 10 and 12 months of age were used in the experiment in **Supplementary Fig. 9**. 6-week-old Nu/J mice for teratoma assays were obtained from The Jackson Laboratory (Cat #00219).

### Doxycycline administration

Doxycycline (Sigma) was added in the drinking water at 1 mg/ml, unless otherwise specified. To improve taste, water was supplemented with 7.5% sucrose. Mice were let to drink water ab libitum, which was kept in amber water bottles to protect the drug from degradation. Fresh doxycycline water was replaced weekly for the duration of the experiment. Doxycycline water was administered continuously for 7, 12 or 18 days, or for 6 cycles of 2-days-doxycycline-ON, 5-days-doxycycline-OFF, as specified in each study.

### Fluorescence activated cell sorting (FACS) of cardiac cells

Hearts from Myh6-Cre^+^Col1a1^OKSM^ (n=5) or Myh6-Cre^−^Col1a1^WT^ mice (n=2) were digested into single cell suspensions following a Langendorff-free method as previously described ^27^ and filtered through a 100 µm strainer (Corning). Cardiac single cell suspensions were stained with cell-specific antibodies (**Supplementary Table 1**) at the concentrations recommended by the manufacturers and sorted on a MoFlo Astrios with a 200 µm nozzle at 9psi (Beckman Coulter) at the Bauer Core Facility of Harvard University.

### Sorting of cardiac cell types by centrifugation and magnetic separation

Myh6-Cre^+^Col1a1^OKSM^ mice (n=5) were administered 1 mg/ml doxycycline in the drinking water for 7 days, followed by euthanasia and digestion of the heart as previously described ^27^. Cardiac myocytes were isolated by two rounds of centrifugation (5 min each) at 30 g. The non-myocyte fraction was then subjected to another round of centrifugation at 30 g to remove any remaining cardiomyocytes. Cardiac endothelial cells were subsequently isolated from the rest of non-myocytes by magnetic cell sorting using a neonatal cardiac endothelial cell isolation kit (Miltenyi) and following the manufacturer’s instructions. The remaining non-myocytes are likely enriched in cardiac fibroblasts and some immune cells but were not sorted to purity. Isolated cell types were further lysed in 1 ml Purezol (Biorad) to proceed with RNA extraction.

### Gene expression by real-time RT-qPCR

Total RNA was extracted from sorted cardiac cells or from cardiac tissues (atria or ventricle) using the Aurum total RNA fatty and fibrous tissues kit (Biorad) and following the manufacturer’s instructions. Tissue samples were homogenized in 1 ml Purezol (Biorad) on a gentleMACS® Octo dissociator (Miltenyi) using M tubes (Miltenyi). RNA quantity and quality were determined by spectrophotometry (Nanodrop). 0.1 µg RNA (sorted cells) or 1 µg RNA (tissues) was used to prepare cDNA with the iScript™ cDNA synthesis kit (Biorad) with the following protocol: 5°C for 5 minutes (min), 46°C for 20 min, 95°C for 1 min. iTaq Universal SYBR green (Biorad) was used to perform real-time qPCR reaction on a CFX96 real-time PCR system (Biorad), following the manufacturer’s protocol: 95°C for 30 seconds (s), followed by 40 cycles of 95°C for 5 s, 60°C for 30 s. A melt curve from 65°C to 95°C was performed at the end of the run to ensure amplification of a single product. *Mapk1* and *Rps13*, identified as the most stable reference genes during cardiac reprogramming in a previous report^17^, were used for data normalization by the Livak method^50^. The list of primers used in this study is found in **Supplementary Table 2**. Real-time RT-qPCR studies were performed in compliance with the Minimum Information for Publication of Quantitative Real-Time PCR Experiments (MIQE) Guidelines^51^. For statistical analysis, ΔCt values were used.

### *In vitro* reprogramming experiments

Cardiomyocytes from Myh6-Cre^+^Col1a1^OKSM^ and Myh6-Cre^−^Col1a1^OKSM^ mice were isolated following a Langendorff-free procedure followed by gravity sedimentation as previously described ^27^. Cells were initially plated on laminin-coated dishes and plating medium and, after 1 h, switched to cardiomyocyte culture medium. The composition of all cell culture media used in this study is summarized in **Supplementary Table 3**. The next day, cultures were switched to embryonic stem cell (ES) culture medium with or without doxycycline (2 μg/ml). This was considered day 0 of the reprogramming experiments. Culture medium with fresh doxycycline was then replenished every other day (50% volume replenished each time). CM cultures were split 1:6 on Matrigel-coated dishes on day 14, and iPSC-like colonies appeared in the doxycycline conditions only. Colonies were expanded on Matrigel coated dishes for two further weeks, at which point doxycycline was progressively weaned off the culture medium for one week. At that point, CM-iPSC colonies were either cryopreserved or used for downstream analysis.

### Immunocytochemistry of CM-iPSC colonies

CM-iPSC colonies cultured on Matrigel-coated surfaces were washed 3 times with PBS, fixed with ice-cold methanol and stained with primary and secondary antibodies (**Supplementary Table 4)** following a standard protocol. Confocal microscopy images (Z stacks) were taken with an LSM 900 microscope (Zeiss) at 20X magnification and processed with Image J software.

### Teratoma assay (CM-iPSC colonies)

CM-iPSC colonies were dissociated into single cell suspensions with trypsin and washed twice by centrifugation with PBS. 10^6^ cells were injected in a 1:1 mix of Matrigel and ice-cold PBS per flank of a Nu/J mice. Teratomas were allowed to form for 4 weeks, and subsequently dissected and fixed in 10% formalin prior to histological analysis by hematoxylin and eosin (H&E) staining and GFP immunohistochemistry following routine protocols, which was performed at iHisto (Boston, USA). Details of antibodies used in staining studies in this work are shown in **Supplementary Table 4**.

### Transmission electron microscopy (TEM)

Myh6-Cre^+^Col1a1^OKSM^ mice were given drinking water with 1 mg/ml doxycycline for 3, 7, 12 or 18 days, or otherwise provided with unadulterated drinking water (n=2 mice/group). At the end of the doxycycline treatment, mice were euthanized and 0.5 ml of 1M KCl was injected through the left ventricle to arrest the heart in diastole, followed by perfusion with 1 ml of TEM fixative (2.5% glutaraldehyde and 2% formaldehyde in 0.1M Sodium cacodylate buffer, pH 7.4). Hearts were then cut into 1-2 mm cubes and incubated in fresh fixative for at least 2 hours (h) at room temperature. Samples were processed for TEM imaging at the Electron Microscopy Facility (Harvard Medical School) following standard techniques. 5 random FOV at 3000X and 5 random FOV at 10000X were imaged from each heart sample (n=2 mice per group).

### Teratoma assay

Myh6-Cre^+^Col1a1^OKSM^ (n=7) and Myh6-Cre^+^Col1a1^WT^ (n=4) mice were given drinking water with 1 mg/ml doxycycline for 18 days and euthanized at the end of the treatment. Hearts were digested into a single cell suspension as described above but, for each heart, cells from the ventricles and from the atria were processed separately. The cells were filtered through a 100 µm strainer (Corning) and washed three times by centrifugation (450 g, 5 min) in ice-cold PBS. Cell pellets were finally resuspended in a 100 µl 1:1 mix of ice-cold PBS and reduced-growth factor Matrigel (Corning) and injected subcutaneously in 6-week-old Nu/J mice (#00219, The Jackson Laboratory). The atrial and ventricular cell suspension of each digested heart were injected in the left and right dorsal flank of a Nu/J mouse, respectively. Teratomas were let to develop for approximately four weeks and then processed for H&E and GFP staining as described above.

### Investigation of the fate of *in vivo* reprogrammed cardiomyocytes (generation of teratomas in the heart)

Myh6-Cre^+^Col1a1^OKSM^ (n=8), Myh6-Cre^+^Col1a1^WT^ (n=14) and Myh6-Cre^−^ Col1a1^OKSM^ (n=10) mice were given drinking water with 1 mg/ml doxycycline for 18 days and then returned to unadulterated drinking water. Mice were closely monitored for changes in body weight and for signs of distress. When mice were found dead or with evident signs of distress, hearts were dissected and washed in PBS. Tissue samples were collected and processed for gene expression (stored in RNA later until RNA isolation was performed as described above) and for histological evaluation.

### Cryosectioning and immunofluorescence

Cardiac teratomas were washed in PBS, embedded in OCT and flash-frozen in isopentane (Sigma) pre-cooled in liquid nitrogen. 10 µm thick sections were obtained with a cryostat (Leica) and processed for immunofluorescence following a standard protocol. The antibodies and dilutions used in this study are listed in **Supplementary Table 3**. Images were acquired with an Axio Scan slide scanner (Zeiss) and processed with QuPath open software (version 0.2.3)^52^.

### *In vivo* reprogramming in aged mice

Male mice of Myh6-Cre^+^Col1a1^OKSM^ and Myh6-Cre^+^Col1a1^OKSM^ genotypes were included in this study. Doxycycline administration (1 mg/ml) was initiated at 285 days of age (∼9 months), and the treatment was performed for 6 cycles of 2-days-ON and 5-days-OFF the drug. Systolic heart function was assessed by echocardiography the day before the start of the treatment (t_0_, baseline) and one day before the end of the study (t_f_). Mice were sacrificed on the last day of the ON phase of the 6^th^ cycle (day 322 of age, ∼10.7 months) and their hearts were digested following a Langendorff-free procedure ^27^ with some modifications. In brief, after digestion, the ventricular cell suspension went through 5 min centrifugation at 30 g, to separate cardiomyocytes from non-myocytes. The cell pellet (cardiomyocytes) was resuspended in perfusion buffer and subjected to a second round of centrifugation to increase purity. The supernatants of both rounds of centrifugation (containing non-myocytes) were then centrifuged again at 30 g (5 min) to remove contaminating cardiomyocytes and later spun down at 450 g (5 min) to collect non-myocytes. Purified cardiomyocytes and non-myocytes were then split in two aliquots, one flash-frozen for DNA sequencing studies and one lysed in Purezol (Biorad) to obtain RNA for RT-qPCR studies.

### DNA methylation studies

DNA was isolated using Qiagen DNeasy Blood & Tissue Kit (Qiagen 69506), eluted in 45 µl of water. The eluted DNA samples were then analyzed by Infinium array HorvathMammalMethylChip320. For the epigenetic age analyses, clocks based on the mouse heart and multi-tissue, and universal mammalian clocks adjusted respectively on age relative to maximum lifespan and age of sexual maturity, are applied to the samples.

### Echocardiography

Systolic heart function was evaluated by echocardiography on a Vevo 3100 high-frequency ultrasound imaging system (Fujifilm, VisualSonics). Inhalable anesthesia (Isoflurane) was kept at 2% throughout the measurements and body temperature was maintained at 37°C by use of a heated platform and heating lamp. Heart rate was maintained at >450 beats per minute (bpm). Ejection fraction (EF, %) was calculated using Vevolab Imaging Software (Fujifilm, VisualSonics).

### Statistical analysis

Statistical analysis was performed with IBM SPSS® Statistics software (version 25). The specific tests applied for each study are described in the respective figure legend. In brief, when dealing with multiple comparisons, Levene’s test of homogeneity of variances was first applied. one-way ANOVA and Tukey’s post-hoc test were used when the variances were homogeneous while Welch ANOVA and Games-Howell post-hoc test were used when the variances of the samples were significantly different. When two groups were compared, an independent samples t test was performed.

## DATA AVAILABILITY

DNA methylation data generated in this study will be made available in GEO upon publication. Datasets generated and/or analysed during the current study are available from the corresponding author on reasonable request. This paper does not report original code.

## ACKNOWLEDGEMENTS

We thank Prof Konrad Hochedlinger (Massachusetts General Hospital and Harvard University) for kindly providing B6;129S4-*Col1a1^tm1(tetO–Pou5f1,–Klf4,–Sox2,–Myc)Hoch^*/J mice. Electron microscopy imaging, consultation and processing of samples were performed at the HMS Electron Microscopy Facility. We thank Zachary Niziolek and the Bauer Core Facility at Harvard University for consultation, performance and analysis of FACS experiments, and staff at the Harvard Bioimaging Facility for assistance with imaging experiments. We thank the Harvard Catalyst Biostatistical Consultation program for advice on statistical analysis of our data. We thank Ahmed Ben Romdhane for his contribution to preliminary cell culture studies. IdL wishes to thank Fred Roberts (Fujifilm) and Dr Dr Jessica Garben (Harvard University) for training in the assessment of cardiac function by echocardiography and for helpful discussions. IdL wishes to thank Matthew Pezone, Mason Dacus and Dr Maxence Dellacherie for their help maintaining the reprogrammable mice colony. DM and IdL wish to acknowledge funding from the Wyss Institute for Biologically Inspired Engineering and from the Initiative for Advanced Biomedical Instrumentation (The University of Hong Kong and Harvard University).

## AUTHOR CONTRIBUTIONS

IdL and DM conceived the study. IdL performed all the experiments. BZ and VNG conducted, analyzed and interpreted DNA methylation studies. NEM and MM contributed to reprogramming studies in old mice. TOS contributed to some RT-qPCR experiments and CMT assisted with sample processing for the teratoma study. IdL and DM wrote the manuscript. All co-authors provided their edits to the manuscript.

## SUPPLEMENTARY FIGURES AND TABLES

**Supplementary Fig. 1. Flow cytometry investigation of GFP^+^ cardiac cell populations in Myh6-Cre^−^Col1a1^WT^ mice (n=2).**

**Supplementary Fig. 2. Gene expression in different cardiac cell types sorted after doxycycline administration.**

**Supplementary Fig. 3. Oct3/4, Sox2 and GFP mRNA levels in off-target organs.**

**Supplementary Fig. 4. Reprogramming of adult mouse cardiomyocytes from Myh6-Cre^+^Col1a1^OKSM^ mice into iPSCs *in vitro*.**

**Supplementary Fig. 5. OKSM expression and pluripotency induction requires Myh6-specific Cre recombination, doxycycline administration and presence of the OKSM cassette.**

**Supplementary Fig. 6. OKSM expression in response to doxycycline (dose and time).**

**Supplementary Fig. 7. Changes in gene expression of TET enzymes upon doxycycline administration.**

**Supplementary Fig. 8. TEM of cardiac ventricle tissue of reprogrammable mice upon doxycycline administration.**

**Supplementary Fig. 9. Cardiac morphology and function in Myh6-Cre^−^ and Myh6-Cre^+^ mice over time.**

**Supplementary Fig. 10. Gene expression in cardiac teratomas and ventricular myocardium.**

**Supplementary Fig. 11. Teratoma assay with *in vivo* reprogrammed cardiomyocytes.**

**Supplementary Fig. 12. Gene expression changes in atrial tissue upon doxycycline administration.**

**Supplementary Fig. 13. TEM of cardiac atrial tissue of reprogrammable mice upon doxycycline administration.**

**Supplementary Fig. 14. Doxycycline-induced differences in DNA methylation age in reprogrammable or control mice.**

**Supplementary Fig. 15. Effects on cardiac function of cyclic administration of doxycycline (2-day ON/5-day OFF) in aged Myh6-Cre^−^Col1a1^WT^ mice.**

**Supplementary Table 1. Antibodies used in FACS studies.**

**Supplementary Table 2. List of primer sequences used in RT-qPCR studies in this work.**

**Supplementary Table 3. Composition of cell culture media used in this study.**

**Supplementary Table 4. Antibodies used for immunohistochemistry and immunofluorescence in this study.**

## References

1 Yilmazer, A., de Lazaro, I., Bussy, C. & Kostarelos, K. In vivo cell reprogramming towards pluripotency by virus-free overexpression of defined factors. PLoS One 8, e54754, (2013).

2 Abad, M. et al. Reprogramming in vivo produces teratomas and iPS cells with totipotency features. Nature 502, 340–345, (2013).

3 de Lazaro, I. et al. Non-viral, Tumor-free Induction of Transient Cell Reprogramming in Mouse Skeletal Muscle to Enhance Tissue Regeneration. Mol Ther 27, 59–75, (2019).

4 Polo, Jose M. et al. A Molecular Roadmap of Reprogramming Somatic Cells into iPS Cells. Cell 151, 1617–1632, (2012).

5 Ocampo, A. et al. In Vivo Amelioration of Age-Associated Hallmarks by Partial Reprogramming. Cell 167, 1719–1733 e1712, (2016).

6 Lu, Y. et al. Reprogramming to recover youthful epigenetic information and restore vision. Nature 588, 124–129, (2020).

7 de Lazaro, I. & Kostarelos, K. In vivo cell reprogramming to pluripotency: exploring a novel tool for cell replenishment and tissue regeneration. Biochemical Society Transactions 42, 711–716, (2014).

8 de Lazaro, I., Cossu, G. & Kostarelos, K. Transient transcription factor (OSKM) expression is key towards clinical translation of in vivo cell reprogramming. EMBO Mol Med, (2017).

9 Wang, C. et al. In vivo partial reprogramming of myofibers promotes muscle regeneration by remodeling the stem cell niche. Nature Communications 12, 3094, (2021).

10 Hishida, T. et al. In vivo partial cellular reprogramming enhances liver plasticity and regeneration. Cell Rep 39, 110730, (2022).

11 Doeser, M. C., Scholer, H. R. & Wu, G. Reduction of Fibrosis and Scar Formation by Partial Reprogramming In Vivo. Stem Cells, (2018).

12 Browder, K. C. et al. In vivo partial reprogramming alters age-associated molecular changes during physiological aging in mice. Nat Aging 2, 243–253, (2022).

13 Chondronasiou, D. et al. Multi-omic rejuvenation of naturally aged tissues by a single cycle of transient reprogramming. Aging Cell 21, e13578, (2022).

14 Alle, Q. et al. A single short reprogramming early in life initiates and propagates an epigenetically related mechanism improving fitness and promoting an increased healthy lifespan. Aging Cell 21, e13714, (2022).

15 Seo, J. H. et al. In Situ Pluripotency Factor Expression Promotes Functional Recovery From Cerebral Ischemia. Mol Ther 24, 1538–1549, (2016).

16 Senis, E. et al. AAVvector-mediated in vivo reprogramming into pluripotency. Nat Commun 9, 2651, (2018).

17 Kisby, T. et al. Adenoviral Mediated Delivery of OSKM Factors Induces Partial Reprogramming of Mouse Cardiac Cells In Vivo. Advanced Therapeutics 4, 2000141, (2021).

18 Eschenhagen, T. et al. Cardiomyocyte Regeneration: A Consensus Statement. Circulation 136, 680–686, (2017).

19 Eminli, S. et al. Differentiation stage determines potential of hematopoietic cells for reprogramming into induced pluripotent stem cells. Nature Genetics 41, 968–976, (2009).

20 Polo, J. M. et al. Cell type of origin influences the molecular and functional properties of mouse induced pluripotent stem cells. Nat Biotechnol 28, 848–855, (2010).

21 Tan, K. Y., Eminli, S., Hettmer, S., Hochedlinger, K. & Wagers, A. J. Efficient generation of iPS cells from skeletal muscle stem cells. PLoS One 6, e26406, (2011).

22 Kisby, T., de Lázaro, I., Stylianou, M., Cossu, G. & Kostarelos, K. Transient reprogramming of postnatal cardiomyocytes to a dedifferentiated state. PLOS ONE 16, e0251054, (2021).

23 Chen, Y. et al. Reversible reprogramming of cardiomyocytes to a fetal state drives heart regeneration in mice. Science 373, 1537–1540, (2021).

24 Agah, R. et al. Gene recombination in postmitotic cells. Targeted expression of Cre recombinase provokes cardiac-restricted, site-specific rearrangement in adult ventricular muscle in vivo. J Clin Invest 100, 169–179, (1997).

25 Stadtfeld, M., Maherali, N., Borkent, M. & Hochedlinger, K. A reprogrammable mouse strain from gene-targeted embryonic stem cells. Nat Methods 7, 53–55, (2010).

26 Belteki, G. et al. Conditional and inducible transgene expression in mice through the combinatorial use of Cre-mediated recombination and tetracycline induction. Nucleic Acids Res 33, e51, (2005).

27 Ackers-Johnson, M. et al. A Simplified, Langendorff-Free Method for Concomitant Isolation of Viable Cardiac Myocytes and Nonmyocytes From the Adult Mouse Heart. Circulation research 119, 909–920, (2016).

28 David, L. & Polo, J. M. Phases of reprogramming. Stem Cell Research 12, 754–761, (2014).

29 de Lazaro, I. et al. Generation of induced pluripotent stem cells from virus-free in vivo reprogramming of BALB/c mouse liver cells. Biomaterials 35, 8312–8320, (2014).

30 Huang, W.-Y., Aramburu, J., Douglas, P. S. & Izumo, S. Transgenic expression of green fluorescence protein can cause dilated cardiomyopathy. Nature medicine 6, 482–483, (2000).

31 Pugach, E. K., Richmond, P. A., Azofeifa, J. G., Dowell, R. D. & Leinwand, L. A. Prolonged Cre expression driven by the α-myosin heavy chain promoter can be cardiotoxic. Journal of molecular and cellular cardiology 86, 54–61, (2015).

32 Rehmani, T., Salih, M. & Tuana, B. S. Cardiac-Specific Cre Induces Age-Dependent Dilated Cardiomyopathy (DCM) in Mice. Molecules (Basel, Switzerland) 24, (2019).

33 Horvath, S. DNA methylation age of human tissues and cell types. Genome Biol 14, R115, (2013).

34 Gladyshev, V. N. The Ground Zero of Organismal Life and Aging. Trends Mol Med 27, 11–19, (2021).

35 Bell, C. G. et al. DNA methylation aging clocks: challenges and recommendations. Genome Biology 20, 249, (2019).

36 Horvath, S. & Raj, K. DNA methylation-based biomarkers and the epigenetic clock theory of ageing. Nat Rev Genet 19, 371–384, (2018).

37 Lu, A. T., et al. Universal DNA methylation age across mammalian tissues. bioRxiv, 2021.2001.2018.426733, (2022).

38 Arneson, A. et al. A mammalian methylation array for profiling methylation levels at conserved sequences. Nature Communications 13, 783, (2022).

39 Mozhui, K. et al. Genetic loci and metabolic states associated with murine epigenetic aging. eLife 11, e75244, (2022).

40 Wesselschmidt, R. L. The teratoma assay: an in vivo assessment of pluripotency. Methods in molecular biology (Clifton, N.J.) 767, 231–241, (2011).

41 Morkin, E. Control of cardiac myosin heavy chain gene expression. Microscopy research and technique 50, 522–531, (2000).

42 De la Rossa, A. et al. In vivo reprogramming of circuit connectivity in postmitotic neocortical neurons. Nat Neurosci 16, 193–200, (2013).

43 Rouaux, C. & Arlotta, P. Direct lineage reprogramming of post-mitotic callosal neurons into corticofugal neurons in vivo. Nat Cell Biol 15, 214–221, (2013).

44 Kim, J. et al. Reprogramming of postnatal neurons into induced pluripotent stem cells by defined factors. Stem cells (Dayton, Ohio) 29, 992–1000, (2011).

45 Sarkar, T. J. et al. Transient non-integrative expression of nuclear reprogramming factors promotes multifaceted amelioration of aging in human cells. Nat Commun 11, 1545, (2020).

46 Shapiro, S. D. et al. Cyclin A2 induces cardiac regeneration after myocardial infarction through cytokinesis of adult cardiomyocytes. Sci Transl Med 6, 224ra227, (2014).

47 D’Uva, G. et al. ERBB2 triggers mammalian heart regeneration by promoting cardiomyocyte dedifferentiation and proliferation. Nature Cell Biology 17, 627–638, (2015).

48 Aguirre, A. et al. In vivo activation of a conserved microRNA program induces mammalian heart regeneration. Cell Stem Cell 15, 589–604, (2014).

49 Kim, Y., Jeong, J. & Choi, D. Small-molecule-mediated reprogramming: a silver lining for regenerative medicine. Experimental & Molecular Medicine 52, 213–226, (2020).

50 Livak, K. J. & Schmittgen, T. D. Analysis of relative gene expression data using real-time quantitative PCR and the 2(-Delta Delta C(T)) Method. Methods 25, 402–408, (2001).

51 Bustin, S. A. et al. The MIQE guidelines: minimum information for publication of quantitative real-time PCR experiments. Clin Chem 55, 611–622, (2009).

52 Bankhead, P. et al. QuPath: Open source software for digital pathology image analysis. Scientific Reports 7, 16878, (2017).

