## Supplementary Information for "In vivo reprogramming and epigenetic rejuvenation of adult cardiomyocytes ameliorate heart failure in mice"

### SUPPLEMENTARY FIGURES

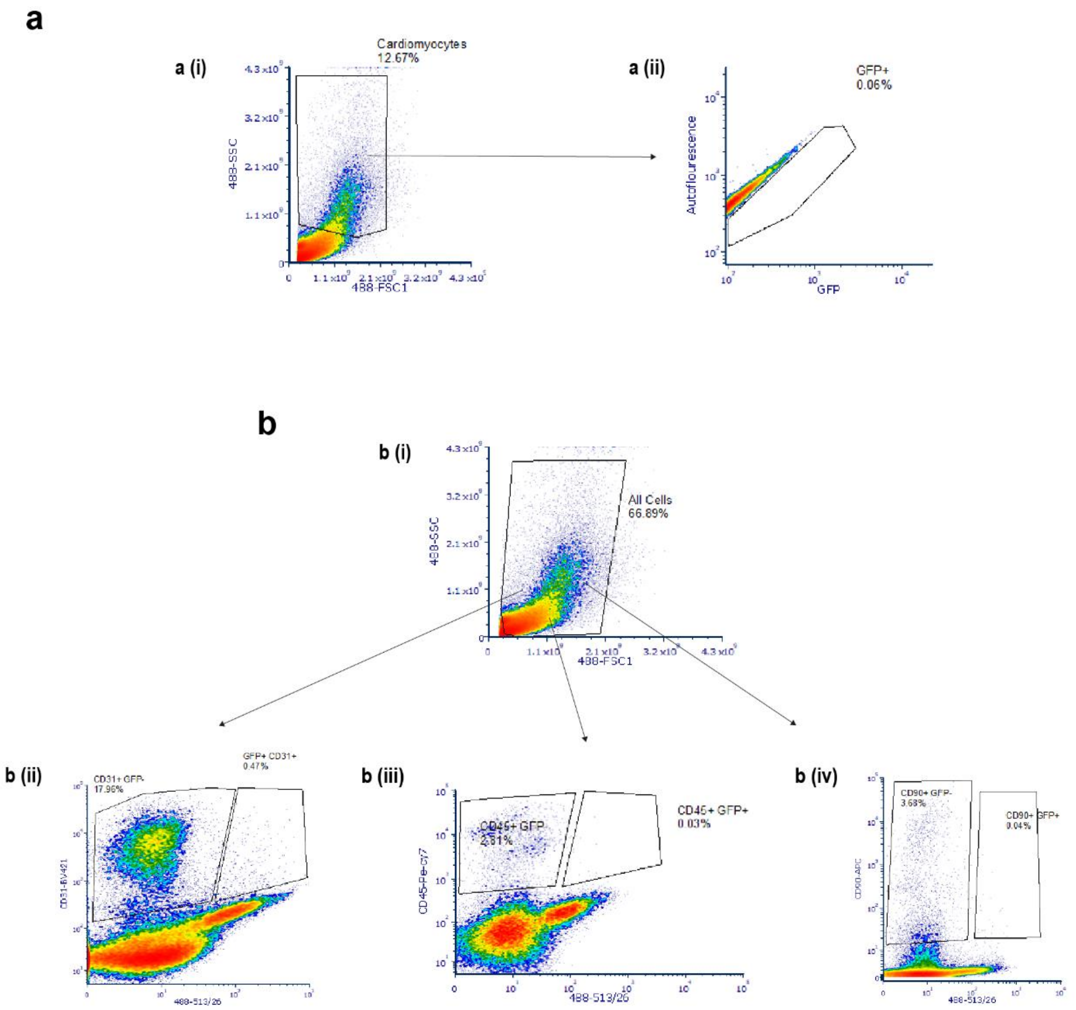


**Figure S1. Flow cytometry investigation of GFP$^{+}$ cardiac cell populations in Myh6-Cre$^{-}$ Col1a1$^{WT}$ mice (n=2). (a)** Representative FACS sorting plots of GFP$^{+}$ cardiomyocytes. **(ai)** Shows sorting of the cardiomyocytes based on their larger size and **(aii)** shows the population of GFP$^{+}$ cells within the cardiomyocyte fraction. **(b)** Sorting of GFP$^{+}$ cells from cardiac cell populations other than cardiomyocytes. **(bi)** Shows gating on all cardiac cells, **(bii)** CD31$^{+}$ endothelial cells, **(biii)** CD45$^{+}$ leukocytes and **(biv)** CD90$^{+}$ fibroblasts.

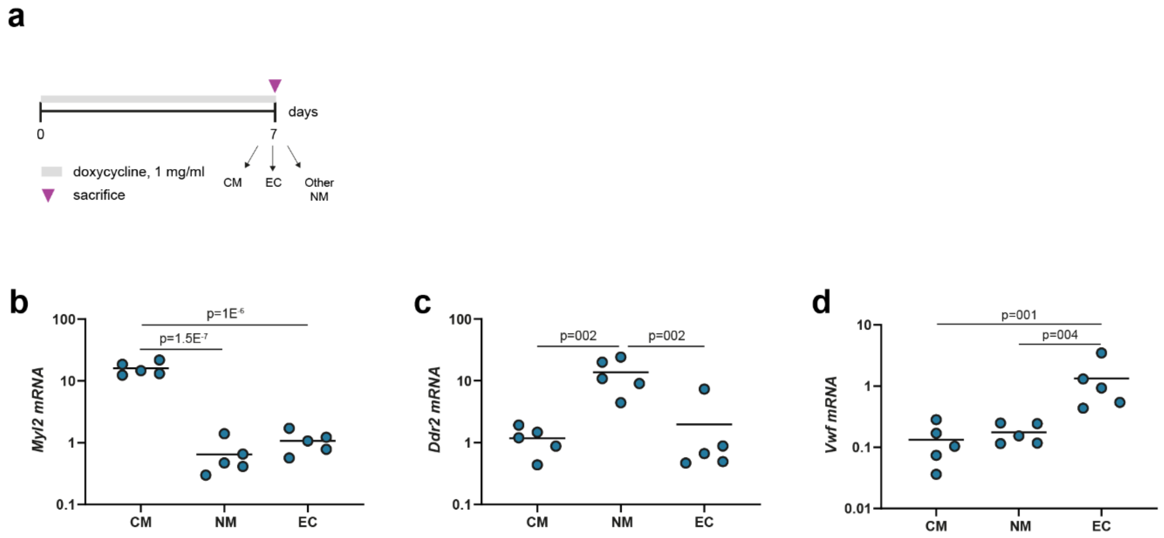


**Figure S2. Gene expression in different cardiac cell types sorted after doxycycline administration. (a)** Schematic representation of the experimental design. mRNA levels of cell-type specific genes including **(b)** *Myl2*, cardiomyocyte-specific; **(c)** *Ddr2*, cardiac fibroblast-specific and **(d)** *Vwf*, endothelial cell-specific, in different cardiac cell types after 7 days of doxycycline administration. CM: cardiomyocyte, NM: non-myocyte, EC: endothelial cell. Gene expression levels were normalized to those in cardiomyocytes. Individual data points represent the fold change ($2^{-\Delta\Delta Ct}$) of each replicate (n=5 mice/group). Statistical analysis was performed by one-way ANOVA and Tukey's post-hoc test.

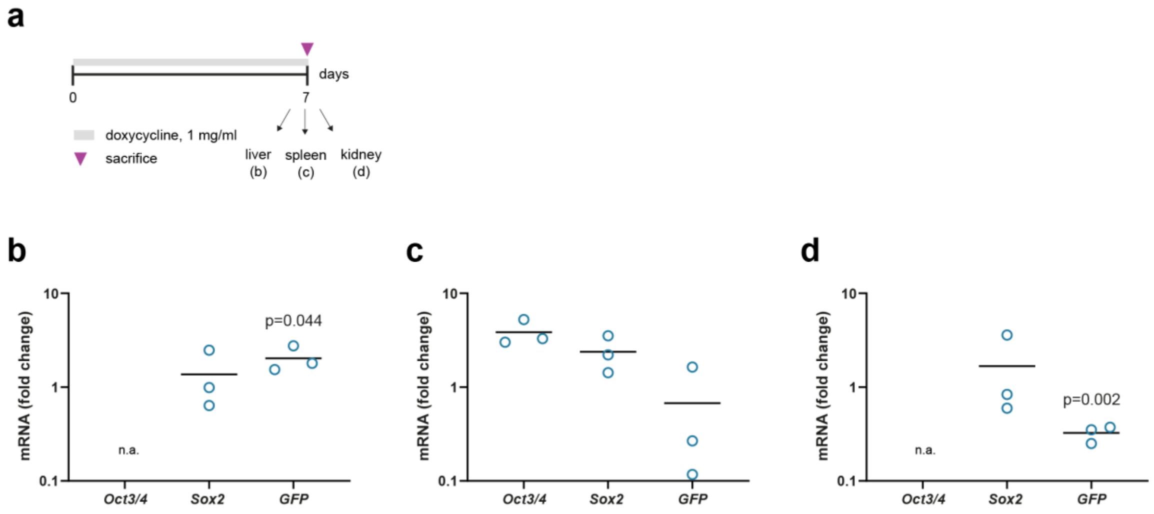


**Figure S3. *Oct3/4*, *Sox2* and *GFP* mRNA levels in off-target organs. (a)** Schematic representation of the experimental design. Gene expression in liver **(b)**, spleen **(c)** and kidney **(d)** tissues of Myh6-Cre$^{+}$Col1a1$^{OKSM}$ mice treated with doxycycline (1 mg/ml) in the drinking water for 7 days. Relative expression was normalized to that of control Myh6-Cre$^{-}$Col1a1$^{OKSM}$ mice under the same treatment. Individual data points represent the fold change ($2^{-\Delta\Delta Ct}$) of each replicate (n=3 mice/group). n.a.: no amplification. Statistical analysis was performed by one-way ANOVA. P values refer to statistical significance in relevance to the control group.

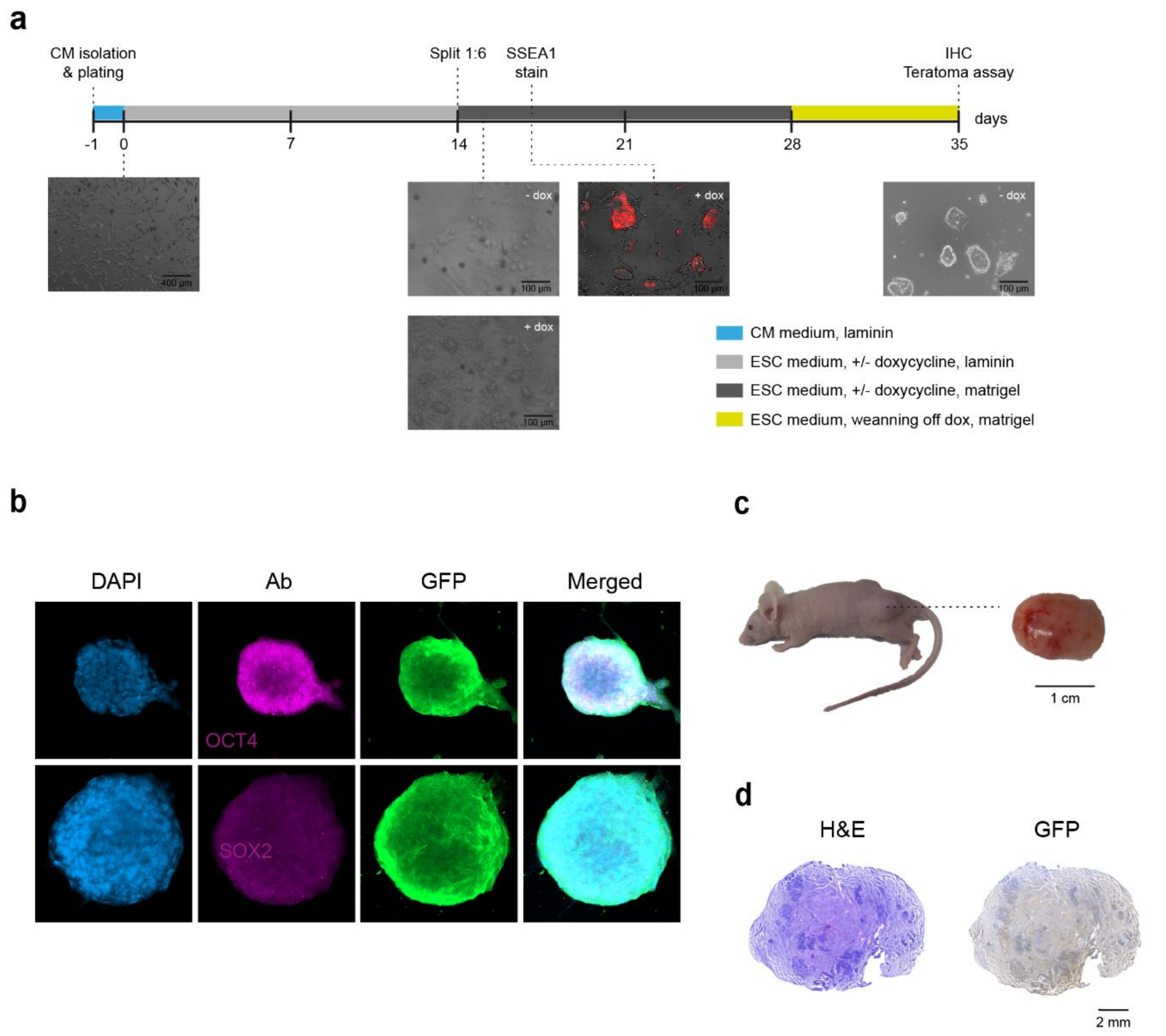


**Figure S4. Reprogramming of adult mouse cardiomyocytes from Myh6-Cre$^{+}$Col1a1$^{OKSM}$ mice into iPSCs *in vitro*. (a)** Schematic representation of the cell culture protocol followed to obtain CM-iPSCs. No colonies appeared in the absence of doxycycline. **(b)** Immunocytochemistry of CM-iPSC colonies. Images were taken at 20X magnification, scale bar = 50 µm. **(c)** Injection of CM-iPSCs into immunodeficient Nu/J mice (teratoma assay) resulted in significant tumor masses. **(d)** H&E and GFP immunostaining of tumors confirmed teratoma identity (trilineage contribution) and cardiomyocyte origin (GFP positive staining).

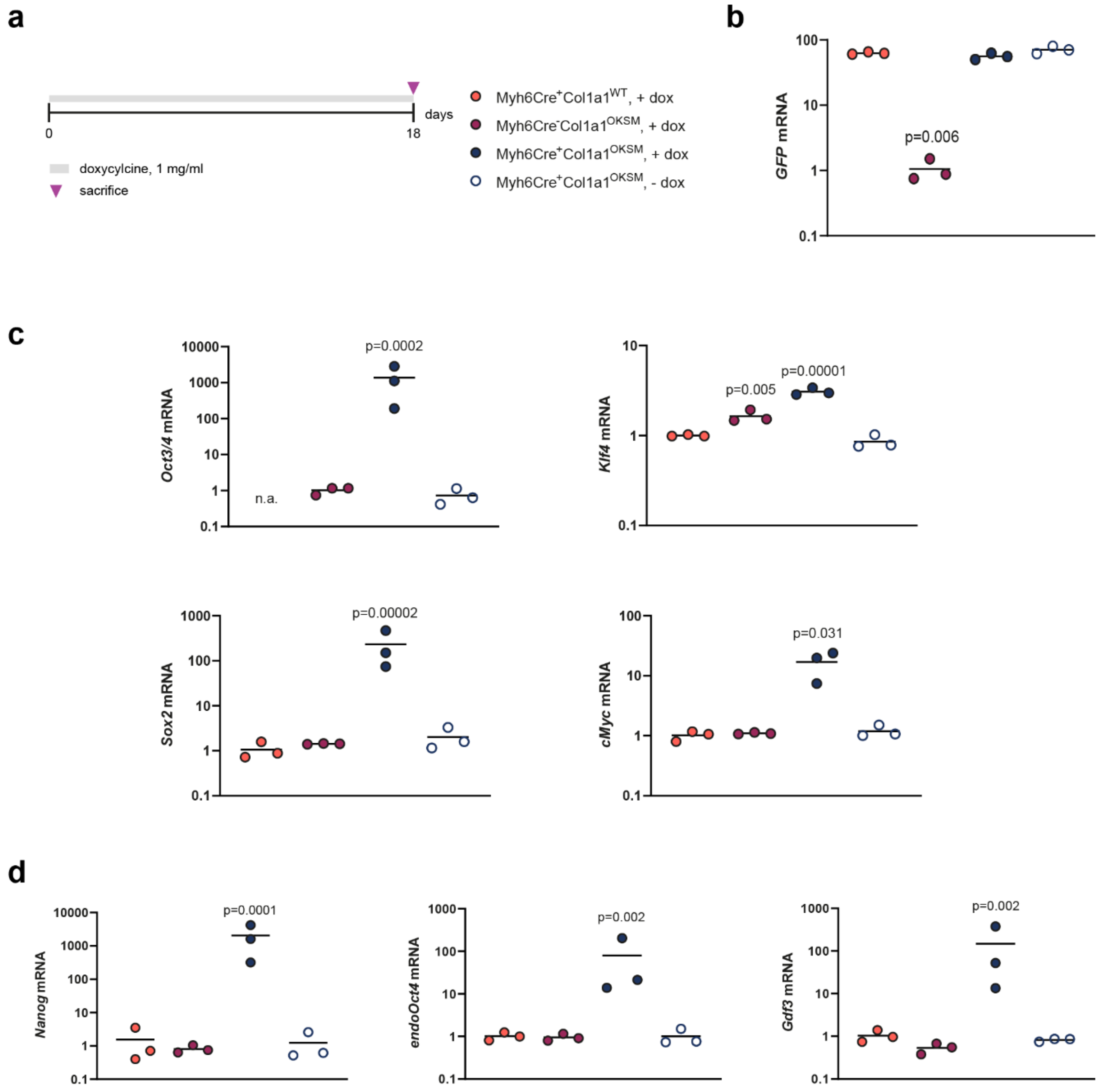


**Figure S5. OKSM expression and pluripotency induction requires Myh6-specific Cre recombination, doxycycline administration and presence of the OKSM cassette. (a)** Schematic representation of the experimental design. mRNA levels of GFP **(b)**, OKSM reprogramming factors **(c)** and endogenous pluripotency genes **(d)** in mice of different genotypes included in the study. GFP levels are normalized to those of Myh6-Cre$^{-}$ Col1a1$^{OKSM}$ mice. *Oct3/4* levels are normalized to those of reprogrammable (Myh6-Cre$^{+}$Col1a1$^{OKSM}$) mice (day 7, 1 mg/ml doxycycline). All other genes are normalized to those of Myh6-Cre$^{+}$Col1a1$^{WT}$ mice. Individual data points represent the fold change ($2^{-\Delta\Delta Ct}$) of each replicate (n=3 mice/group). Statistical analysis was performed by one-way ANOVA and Tukey's post-hoc test. P values refer to statistical significance in relevance to the control group.

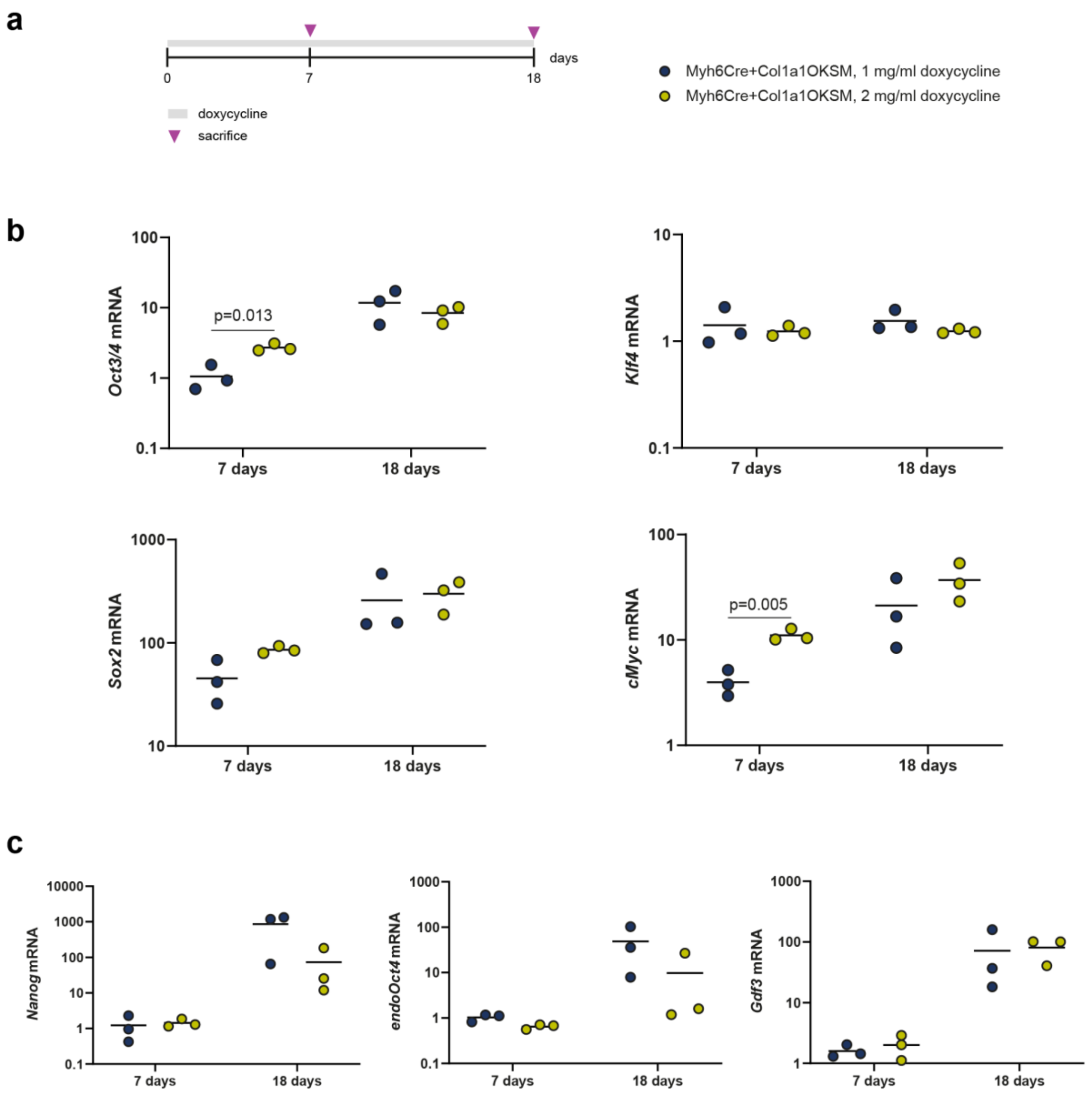


**Figure S6. OKSM expression in response to doxycycline (dose and time). (a)** Schematic representation of the experimental design. **(b)** mRNA expression of reprogramming factors (*Oct3/4*, *Klf4*, *Sox2* and *cMyc*), varying doxycycline dose and time. **(c)** mRNA expression of endogenous pluripotency genes (*Nanog*, *endoOct4* and *Gdf3*), varying doxycycline dose and time. *Oct3/4* gene expression levels are normalized to those of Myh6-Cre$^{+}$Col1a1$^{OKSM}$ mice (7 days doxycycline, 1 mg/ml dose). Data from all other genes are normalized to Myh6-Cre$^{+}$Col1a1$^{WT}$ controls. Individual data points represent the fold change ($2^{-\Delta\Delta Ct}$) of each replicate (n=3 mice/group). Statistical analysis was performed by one-way ANOVA and Tukey's post-hoc test. P vales in the figure represent statistical significance between 1 mg/ml and 2 mg/ml doxycycline treatment groups.

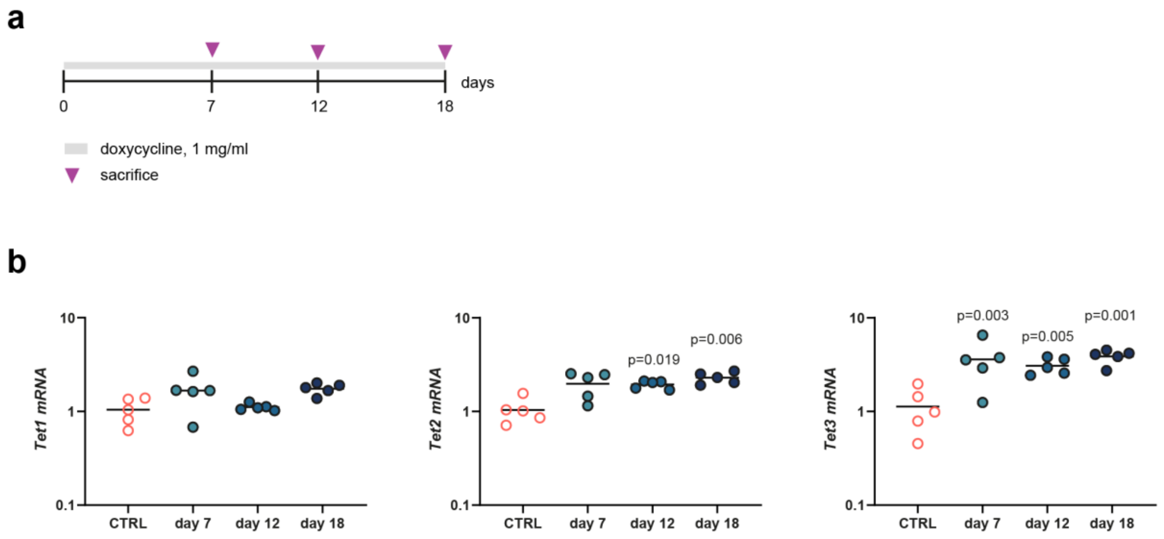


**Figure S7. Changes in gene expression of TET enzymes upon doxycycline administration. (a)** Schematic of experimental design. **(b)** mRNA levels of *Tet1*, *Tet2* and *Tet3* enzymes. Gene expression was normalized to that of control (Myh6-Cre$^{+}$Col1a1$^{WT}$) mice. Individual data points represent the fold change ($2^{\wedge-\Delta\Delta Ct}$) of each replicate (n=5 mice/group). Statistical analysis was performed by one-way ANOVA and Tukey's post-hoc test. P values refer to statistical significance in relevance to the control group.

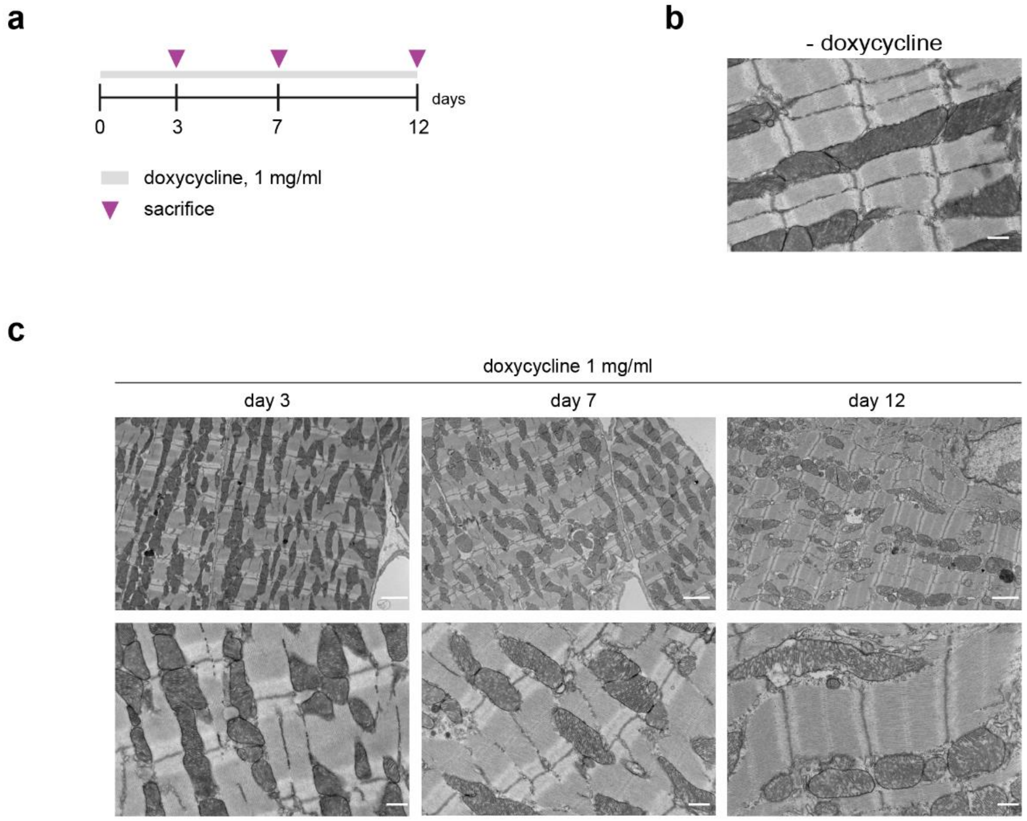


**Figure S8. TEM of cardiac ventricle tissue of reprogrammable mice upon doxycycline administration. (a)** Schematic representation of the experimental design. **(b)** Representative TEM micrograph of cardiac ventricle tissue from reprogrammable mice administered with unadulterated drinking water, Scale bar denotes 500 nm. **(c)** Representative TEM micrographs of cardiac ventricle tissue from reprogrammable mice administered with 1 mg/ml doxycycline in the drinking water for 3, 7 or 12 days**.** Scale bars represent 2 µm on top panel and 500 nm on bottom panel. The figure shows representative images from 5 random FOV and n=2 mice/group.

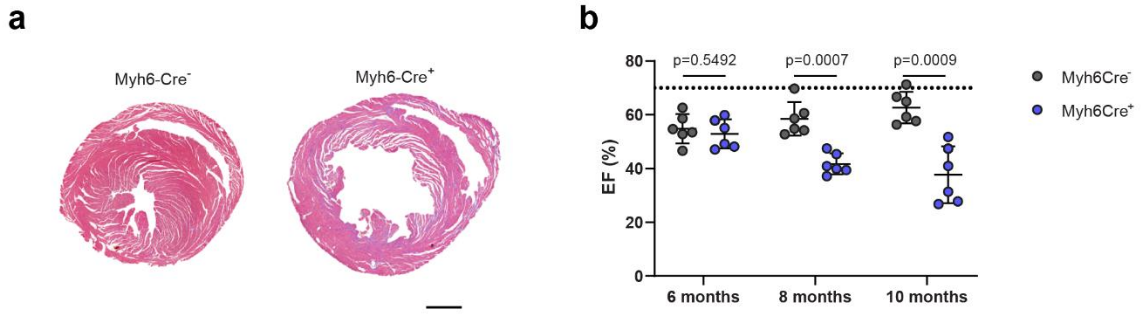


**Figure S9. Cardiac morphology and function in Myh6-Cre⁻ and Myh6-Cre⁺ mice over time. (a)** Representative images of H&E staining of ventricular cross-sections from 1-year-old Myh6-Cre⁻ (left, n=4 mice) and Myh6-Cre⁺ mice (right, n=5 mice). Scale bar = 1 mm. **(b)** Ejection fraction values in Myh6-Cre⁻ and Myh6-Cre⁺ mice at 6, 8 and 10 months of age (n=6). Statistical analysis was performed by independent samples t tests and p values refer to differences between Myh6-Cre⁻ and Myh6-Cre⁺ mice at each age group.

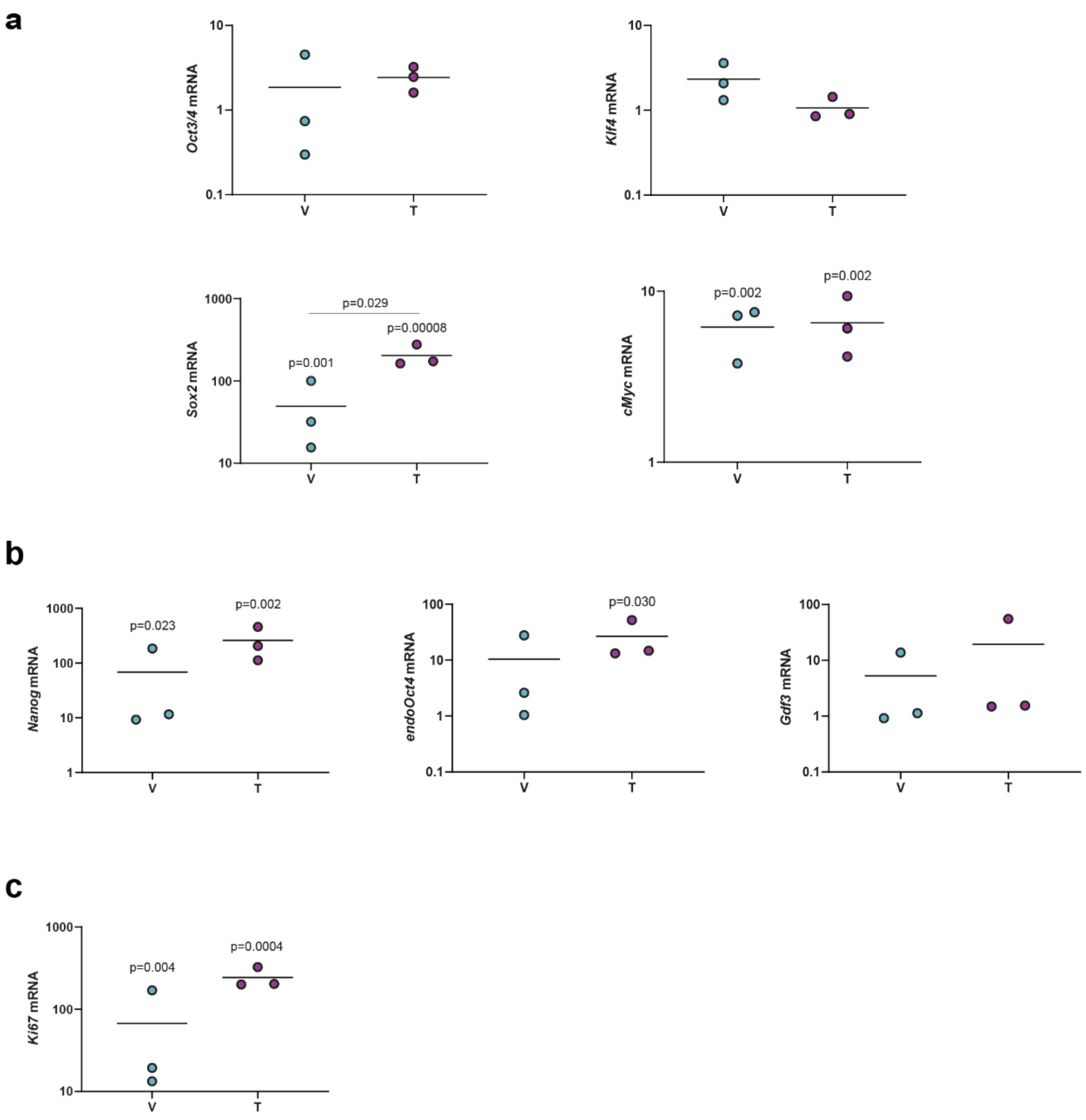


**Figure S10. Gene expression in cardiac teratomas and ventricular myocardium.** Gene expression levels of OKSM reprogramming factors **(a)**, endogenous pluripotency genes **(b)** and *Ki67* **(c)** in cardiac teratomas (T) and ventricular tissue (V) of reprogrammable (Myh6-Cre$^{+}$Col1a1$^{OKSM}$) mice. *Oct3/4* gene expression levels are normalized to those of the ventricle tissue of reprogrammable mice. All other genes are normalized to Myh6-Cre$^{+}$Col1a1$^{WT}$ controls. Individual data points represent the fold change (2^$^{-\Delta\Delta Ct}$) of each replicate (n=3 mice/group). Statistical analysis was performed by one-way ANOVA and Tukey's post-hoc test. P values in the graphs refer to statistical significance in relevance to the control group or between ventricle and teratoma (in *Sox2*, represented with a line between the two groups).

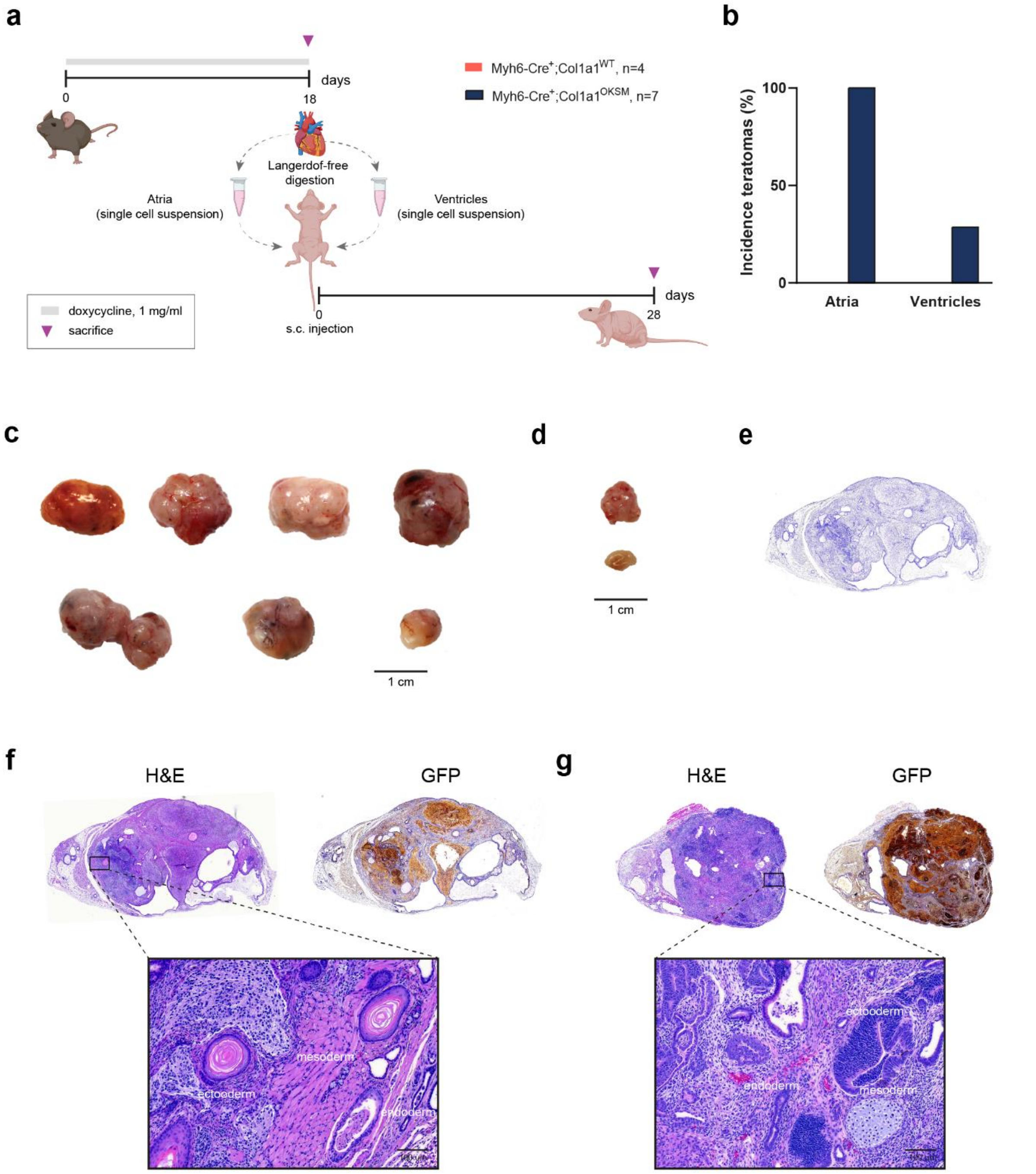


**Figure S11. Teratoma assay with in vivo reprogrammed cardiomyocytes. (a)** Schematic representation depicting the workflow of the teratoma assay. Partially created with BioRender.com **(b)** Number of teratomas generated from atrial and ventricular single cell suspensions from reprogrammable (Myh6-Cre$^{+}$Col1a1$^{OKSM}$) and non-reprogrammable (Myh6-Cre$^{+}$Col1a1$^{OKSM}$) mice administered with doxycycline for 18 days. Images of atrial **(c)** and ventricular **(d)** teratomas 4 weeks after s.c. injection of reprogrammed cardiac cells. Composite images have been created with individual teratoma images maintained at the same scale. Scale bars represent 1 cm. **(e)** Negative control of GFP staining (no primary antibody) in a teratoma section (atrial origin). **(f)** Representative H&E and GFP staining of a teratoma of atrial origin confirms trilineage contribution (high magnification insert, scale bar represents 100 µm) and cardiomyocyte origin (GFP positive staining). **(g)** Representative H&E and GFP staining of a teratoma of ventricular origin confirms trilineage contribution (high magnification insert, scale bar represents 100 µm) and cardiomyocyte origin (GFP positive staining). Figure shows representative images, but trilineage contribution and GFP positive staining were confirmed in all teratomas.

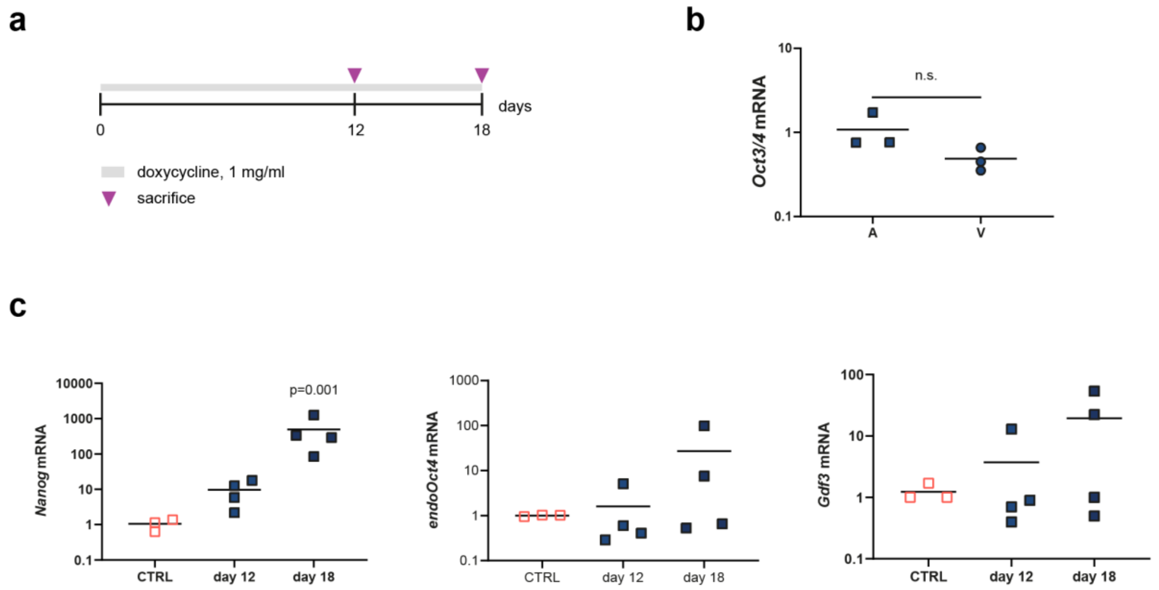


**Figure S12. Gene expression changes in atrial tissue upon doxycycline administration. (a)** Schematic representation of the experimental design. **(b)** *Oct3/4* mRNA levels on atrial and ventricular tissue after 12 days of doxycycline administration. Expression levels were normalized to that of the ventricle. Individual data points represent the fold change ($2^{\wedge\text{-}\Delta\Delta Ct}$) of each replicate (n=3 mice/group). Statistical analysis was performed by one-way ANOVA. n.s.: not significant. **(b)** Expression of endogenous pluripotency genes in atrial tissue 12 and 18 days after the start of doxycycline treatment. Gene expression levels were normalized to those of control (Myh6-Cre$^{+}$Col1a1$^{WT}$) mice. Individual data points represent the fold change ($2^{\wedge\text{-}\Delta\Delta Ct}$) of each replicate (n=3-4 mice/group). Statistical analysis was performed by one-way ANOVA and Tukey's post-hoc test. P values refer to statistical significance in relevance to the control group.

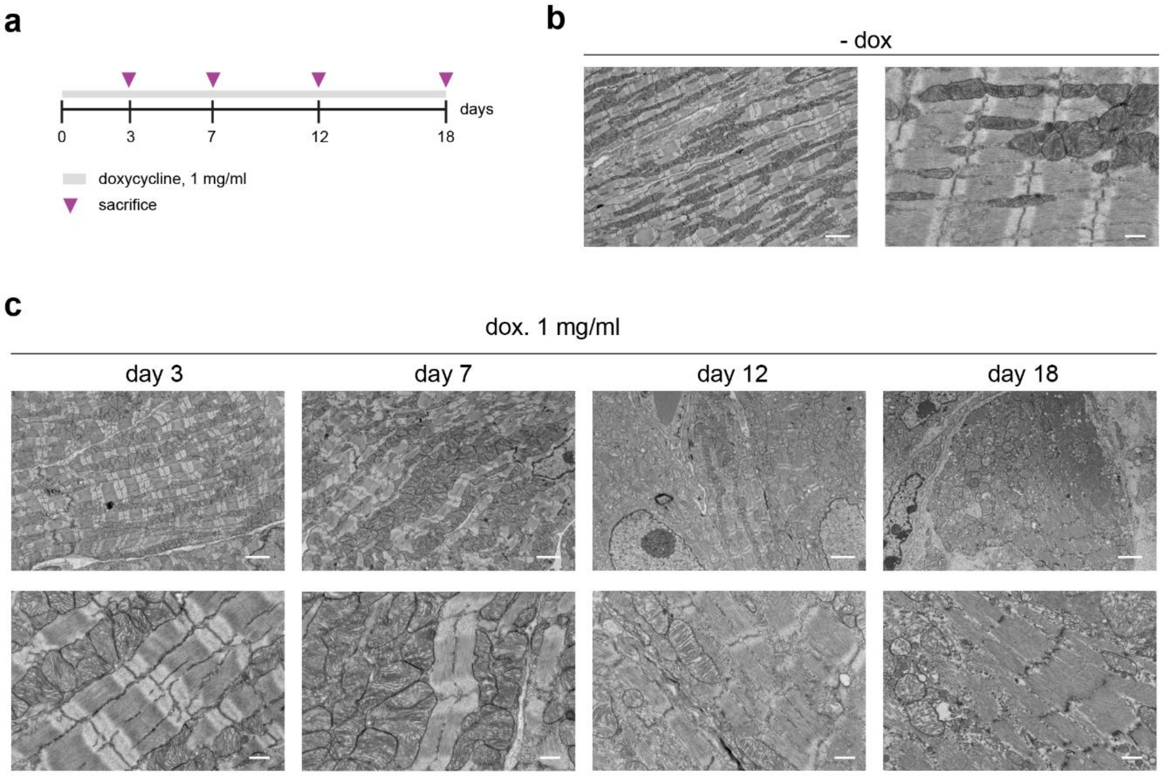


**Figure S13. TEM of cardiac atrial tissue of reprogrammable mice upon doxycycline administration. (a)** Schematic representation of the experimental design. **(b)** TEM micrographs of atrial tissue of reprogrammable mice administered with unadulterated water (without doxycycline). Figure shows representative images from 5 random FOV and n=2 mice. TEM scale bars represent 2 µm (right) and 500 nm (left). **(c)** Representative TEM micrographs of cardiac atrial tissue from reprogrammable mice administered with 1 mg/ml doxycycline in the drinking water for 3, 7, 12 and 18 days**.** Scale bars represent 2 µm on top panel and 500 nm on bottom panel. The figure shows representative images from 5 random FOV and n=2 mice/group.

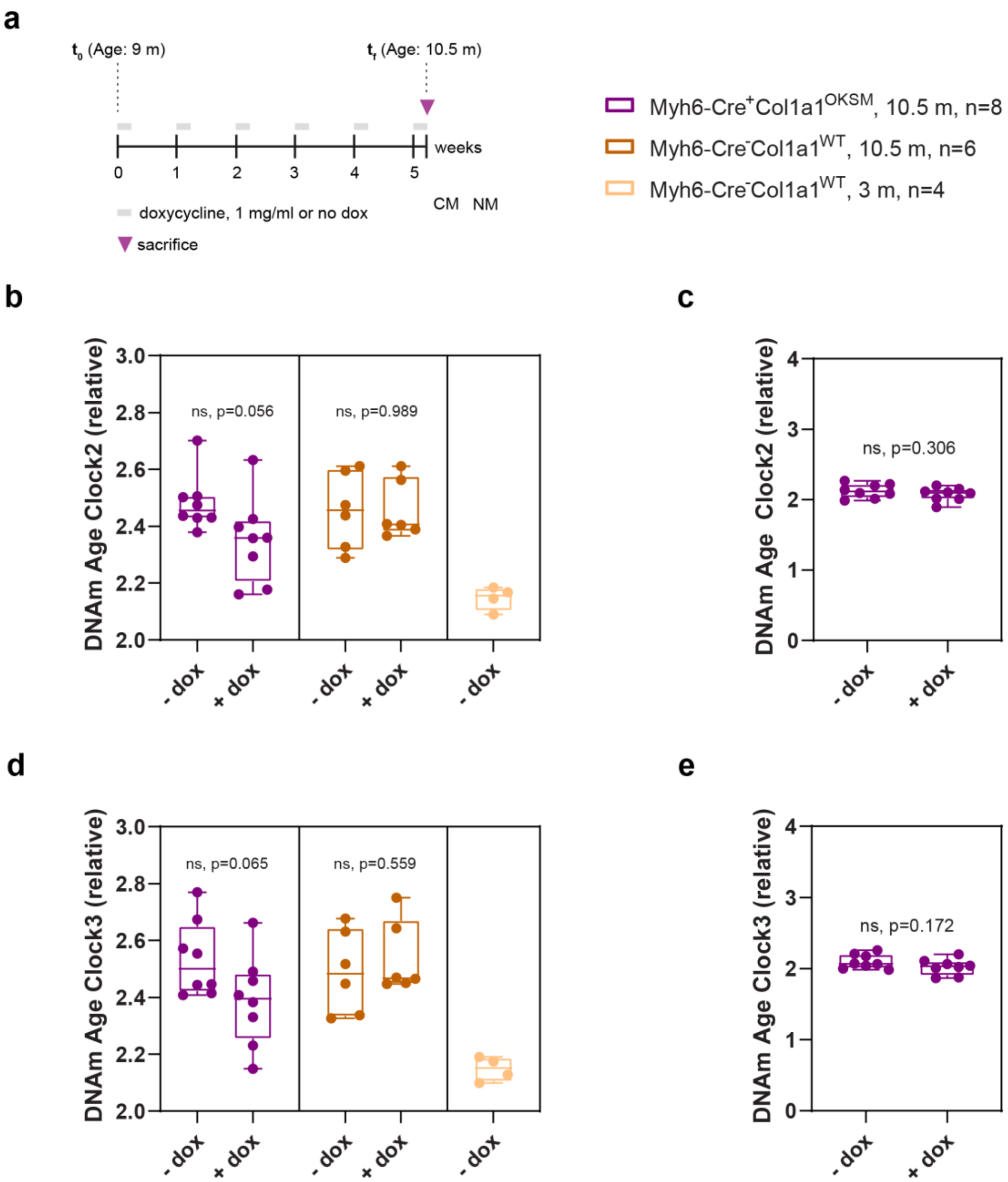


**Figure S14. Doxycycline-induced differences in DNA methylation age in reprogrammable or control mice. (a)** Schematic representation of the experimental design. **(b)** Pan-tissue epigenetic "Clock2" applied to the DNA methylation pattern of cardiomyocytes. **(c)** Pan-tissue epigenetic "Clock2" applied to the DNA methylation pattern of non-myocytes. **(d)** Pan-tissue epigenetic "Clock3" applied to the DNA methylation pattern of cardiomyocytes. **(e)** Pan-tissue epigenetic "Clock3" applied to the DNA methylation pattern of non-myocytes. Statistical differences were calculated by independent samples t tests. P values refer to differences between each treated (+doxycycline) and untreated (-doxycycline) group. ns: not significant.

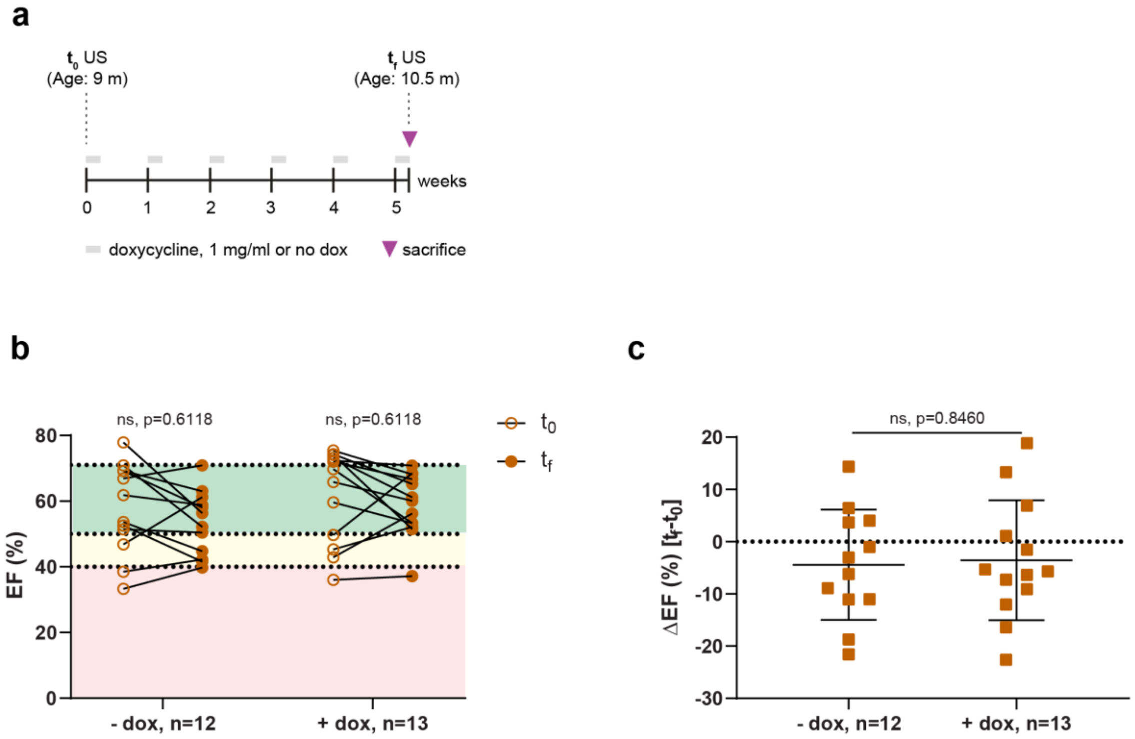


**Figure S15. Effects on cardiac function of *c*yclic administration of doxycycline (2-day ON/5-day OFF) in aged Myh6-Cre$^{-}$Col1a1$^{WT}$ mice. (a)** Schematic representation of experimental design. Doxycycline treatment (6 cycles 2ON/5OFF) was initiated when mice were 9 m of age and systolic heart function was assessed via echocardiography at the start ($t_0$) and end of the study ($t_f$). Ejection fraction (EF) values at $t_0$ and $t_f$ are shown in **(b)** and expressed as the ΔEF ($t_f$-$t_0$) in **(c)**. Statistical analysis was performed by independent samples t Test. P values represent differences between the $t_f$ and $t_0$ (in b) and between the ΔEF of the untreated and treated groups (in c).

### SUPPLEMENTARY TABLES

***Supplementary Table 1. Antibodies used in FACS studies.***

| Cell type | Antigen target | Fluorophore | Company (reference) |
|---|---|---|---|
| Cardiac fibroblasts | CD90 | APC | Thermofisher (A1427) |
| Endothelial cells | CD31 | BV421 | Biolegend (102423) |
| Leucocytes | CD45 | PE-Cy7 | Biolegend (103114) |

***Supplementary Table 2. List of primer sequences used in RT-qPCR studies in this work.***
*Primers for Oct3/4, Klf4, Sox2 and cMyc amplify both the transgene and endogenous form, while primers for endoOct4 are specific for the endogenous mRNAs.*

| Gene symbol | Gene ID (NCBI) | Primer Forward | Primer Reverse |
|---|---|---|---|
| *Mapk1* | 26413 | GGTTGTTCCCAAATGCTGACT | CAACTTCAATCCTCTTGTGAGGG |
| *Rps13* | 68052 | CCCAGGTCCGTTTTGTGACT | TCCTCTCAAGGTGCTTTCGG |
| *Myl2* | 17906 | GGACTGAGCCCTGAACCAC | AGCCAAGACTTCCTGTTTATTTGC |
| *Ddr2* | 18214 | CTGATTGGTTGCTTGGTGGC | TTGAACATGCTGGACTCGCT |
| *Vwf* | 22371 | TGGTTGACCTATGCACGACC | GCATTCACCCTGGCTCTTCT |
| *Oct3/4* | 18999 | TGAGAACCTTCAGGAGATATGCAA | CTCAATGCTAGTTCGCTTTCTCTTC |
| *Klf4* | 16600 | CAGTGGTAAGGTTTCTCGCC | GCCACCCACACTTGTGACTA |
| *Sox2* | 20674 | GGTTACCTCTTCCTCCCACTCCAG | TCACATGTGCGACAGGGGCAG |
| *cMyc* | 17869 | CAGAGGAGGAACGAGCTGAAGCGC | TTATGCACCAGAGTTTCGAAGCTGTTCG |
| *GFP* | 25339618 | AAGCTGACCCTGAAGTTCATCTGC | CTTGTAGTTGCCGTCGTCCTTGAA |
| *Nanog* | 71950 | CAGAAAAACCAGTGGTTGAAGACTAG | GCAATGGATGCTGGGATACTC |
| *endoOct4* | 18999 | TCTTTCCACCAGGCCCCCGGCTC | TGCGGGCGGACATGGGGAGATCC |
| *Gdf3* | 14562 | GTTCCAACCTGTGCCTCGCGTCTT | AGCGAGGCATGGAGAGAGCGGAGCAG |
| *Ki67* | 17345 | CTGGTCACCATCAAGCGGAG | CAATACTCCTTCCAAACAGGCAG |
| *Cdh1* | 12550 | CAGCCGGTCTTTGAGGGATT | TGACGATGGTGTAGGCGATG |
| *Epcam* | 17075 | AACAACGATGGGCTGTACGA | CCGTGTCCTTGTCGGTTCTT |
| *Tet1* | 52463 | TCAAGCAATGGACCACTGGG | TCTCCATGAGCTCCCTGACA |
| *Tet2* | 214133 | ACTCCTGGTGAACAAAGTCAGA | CATCCCTGAGAGCTCTTGCC |
| *Tet3* | 194388 | CCGGATTGAGAAGGTCATCTAC | AAGATAACAATCACGGCGTTCT |
| *Myh6* | 17888 | GCCCAGTACCTCCGAAAGTC | GCCTTAACATACTCCTCCTTGTC |
| *cTnT* | 21956 | CAGAGGAGGCCAACGTAGAAG | CTCCATCGGGGATCTTGGGT |

*Supplementary Table 3. Composition of cell culture media used in this study.*

| Culture medium | Component | Concentration | Supplier |
|---|---|---|---|
| **Plating Medium** | M199 | - | Sigma-Aldrich |
| | FBS | 5% | Gibco |
| | BDM | 10 mM | Acros Organics |
| | PS | 1% | Gibco |
| **Cardiomyocyte medium** | M199 | - | Sigma-Aldrich |
| | BSA | 0.1% | Sigma-Aldrich |
| | ITS | 1X | Corning |
| | BDM | 10 mM | Acros Organics |
| | CD lipid | 1X | Gibco |
| | PS | 1% | Gibco |
| **ESC medium** | KO DMEM | - | Gibco |
| | KSR | 15% | Gibco |
| | β-mercaptoethanol | 50 μM | Sigma-Aldrich |
| | GlutaMax | 1X | Gibco |
| | NEAA | 1X | StemCell |
| | PS | 1% | Gibco |
| | LIF | 10 ng/ml | Merk |

M199: medium 199, FBS: fetal bovine serum, BDM: 2,3-Butanedione Monoxide, PS: Penicillin/Streptomycin, BSA: bovine serum albumin, ITS: insulin/transferrin/selenium, CD lipid: chemically defined lipid concentrate, KO DMEM: knock-out Dulbecco's modified Eagle medium, KSR: knock-out serum replacement, NEAA: non-essential aminoacids, LIF: leukemia inhibitory factor.

***Supplementary Table 4. Antibodies used for immunohistochemistry and immunofluorescence in this study.***

| | **Antibody** | **Company (reference)** | **Dilution** |
|---|---|---|---|
| **Primary** | Chicken anti-GFP | Abcam (ab13970) | 1:1000 |
| | Mouse anti-Cardiac Troponin T | Abcam (ab8295) | 1:500 |
| | Rabbit anti-Ki67 | Abcam (ab15580) | 1:500 |
| | Rabbit anti-Oct4 | Abcam (ab19857) | 1:300 |
| | Rabbit anti-Sox2 | Abcam (ab97959) | 1:1000 |
| | Rabbit anti-Nanog | Abcam (ab80892) | 1:200 |
| **Secondary** | Anti-chicken Alexa Fluor 594 | Thermo Scientific (A32759) | 1:500 |
| | Anti-chicken Alexa Fluor 488 | Jackson Immunoresearch (103545155) | 1:250 |
| | Anti-mouse Alexa Fluor 647 | Jackson Immunoresearch (115605166) | 1:250 |
| | Anti-rabbit Alexa Fluor 750 | ThermoScientific (A21039) | 1:500 |
| | Anti-rabbit Alexa Fluor 680 | ThermoScientific (A32734) | 1:500 |
